# Local disorder is associated with enhanced catalysis in a nascent photoswitch

**DOI:** 10.1101/2024.11.26.625553

**Authors:** James W. McCormick, Jerry C. Dinan, Marielle AX Russo, Kevin H. Gardner, Kimberly A. Reynolds

## Abstract

Domain insertion is a common strategy for introducing allosteric regulation in both engineered and evolved systems. In this approach, an “input” domain is covalently fused to an “output” domain with the goal of conferring new regulation. In prior work, we found that insertion of the LOV2 domain at evolutionarily conserved allosteric “hot spots” on the metabolic enzyme Dihydrofolate Reductase (DHFR) could confer modest light regulation of enzymatic activity. However, it was not clear if the newly established regulation was achieved by interdomain allosteric conformational coupling, or if it represented a “simpler” mechanism like coupling of LOV2 light activation to global folding stability of DHFR or light-dependent steric occlusion of the DHFR active site. To better understand how these newly formed domain fusions harness light-inducible disorder in LOV2 for allosteric activation, we biochemically characterized a representative synthetic fusion. We observed that LOV2 photoactivation simultaneously: (1) thermally destabilized the fusion and (2) lowered the DHFR catalytic transition free energy of the lit state relative to the dark state. Light-induced NMR chemical shift changes indicated that photochemically-initiated conformational changes propagated from LOV2 to the active site of DHFR. Moreover, ligand binding at DHFR modified LOV2 chemical shifts, demonstrating bidirectional coupling between domains. Examination of select allostery-tuning mutations found a modest negative correlation between the light-induced change in thermal stability and catalytic activity, suggesting an activity-stability tradeoff. Together our data indicate that a domain fusion event can realize localized conformational coupling between active sites even in the absence of extensive evolutionary optimization.

## INTRODUCTION

Inducible order-to-disorder transitions play a key role in the allosteric regulation of numerous evolved and engineered proteins ^1^. In many cases, modulation of disorder in one protein domain is used to regulate flexibility and activity in another. As one classic example, the tetracycline repressor (TetR) loses affinity for DNA when tetracycline binding at the ligand binding domain reduces the flexibility of key loops in the DNA binding domain ^2^. Likewise, eukaryotic transcription factors frequently make use of locally disordered domains in regulation ^3^. Following from this naturally observed architecture, inserting an “input domain” with an inducible order-disorder transition into an “output domain” of interest has become a frequent strategy for achieving allostery in engineered systems. This approach has yielded a multitude of enzymes and signaling molecules regulated by ligand, light, pH, and temperature ^4–8^.

Here we investigate how inducible disorder promotes catalysis for a synthetic allosteric fusion between the light-sensing LOV2 domain of *Avena sativa* phototropin 1 (LOV2) and the *Escherichia coli* metabolic enzyme dihydrofolate reductase (DHFR). The LOV2 domain is undergoes a light-driven order-to-disorder change in both its native context as well as in several engineered systems ^9–11^. More specifically, blue light absorption drives the photochemical reduction of the flavin mononucleotide (FMN) cofactor and concomitant formation of a covalent linkage between LOV2 cysteine 450 and the FMN isoalloxazine ring. This triggers allosteric conformational changes in the surrounding protein, leading to unfolding of the A’α and Jα helices at the LOV2 N- and C-termini. The N- and C-termini of LOV2 are physically proximal, and insertion of LOV2 into another protein domain has been used to regulate a number of signaling proteins ^6,7^. However, unlike the light-activation seen for the native phototropin 1 kinase protein, engineered systems often use the LOV2 order-disorder transition to disrupt rather than increase catalytic activity. The general rationale is that light-induced local disorder can destabilize or de-populate the active state ^6,8^. There are a handful of engineered LOV2 photo-activatable systems, but these seem to promote activity via domain dissociation or an uncaging event rather than through allosteric regulation ^12,13^.

As such, the DHFR-LOV2 system studied here is relatively unique among engineered proteins in that it gives rise to light-induced allosteric activation. In this system, analysis of amino acid co-evolution was used to guide insertion of LOV2 into a DHFR loop that undergoes conformational changes associated with catalysis (Fig. 1A) ^5,14^. Because LOV2 is inserted between residues 120 and 121 of DHFR, we call the resulting construct DL121 (Fig. 1B). In prior work, we found that LOV2 insertions at specific DHFR surface sites that co-evolve with the active site (termed sector-connected surfaces) were statistically more likely to yield new regulation ^14^. Indeed, while the DL121 fusion shows modest but statistically significant light activation (a 34% increase in *k_cat_*in the light relative to the dark), insertions of LOV2 into the same DHFR loop just a few residues away do not allosterically signal. We underscore that this level of regulation drives a 20% increase in bacterial growth rate in response to light^14^, an effect which could easily drive evolution if allostery is a condition of selection. Moreover, we previously found that structurally disperse, solvent exposed positions can further tune allostery (either enhancing or disrupting the magnitude of the effect). These mutations individually yield small variations in allosteric effect (on the order of 10-50%) but can be combined to generate larger effects ^15^. Thus, DL121 provides a unique opportunity to understand how domain insertion events give rise to new inter-domain regulation that can serve as a substrate for further evolution. To better characterize the allosteric mechanism of the synthetic DL121 fusion and the associated allostery-tuning mutations, we pursued a combination of CD and NMR spectroscopy and Eyring analysis. We confirmed that the LOV2 domain of DL121 does undergo an order to disorder transition (losing alpha helical content) in response to light, as in other systems. LOV2 insertion destabilizes DHFR, reducing catalytic turnover (*k_cat_*) by 21-fold relative to the native enzyme, and bringing the T_m_ of the fusion to a near-physiological temperature in the dark. This raises the possibility that allosteric activation of DHFR catalysis by LOV2 might work by modulating the global folding equilibrium and increasing the fraction of folded enzyme in the lit state. However, experimental measurements of thermal stability showed that light exposure mildly destabilizes DL121. This would thus reduce the fraction folded in the light, in opposition to the observed apparent increase in activity. We then used Eyring analysis to characterize the thermodynamics of catalysis. Eyring analysis revealed that light activation (and the commensurate folding destabilization) is associated with a favorable decrease in the enthalpy of transition state activation, offset by an entropic penalty. Thus, light-induced local unfolding seems to permit a more enthalpically favorable transition state. Allostery-tuning mutations altered this entropy-enthalpy balance. Using solution NMR, we observed that changes in the DHFR active site (ligand binding) affect LOV2 NMR chemical shifts and vice-versa, indicating bi-directional allosteric coupling between the domains. Together our work shows that engineered fusions can go beyond simply using an input domain to disrupt or break the activity of an output domain, and can instead achieve specific conformational coupling between active sites upon an initial unoptimized, covalent domain fusion event.

**Figure 1.**
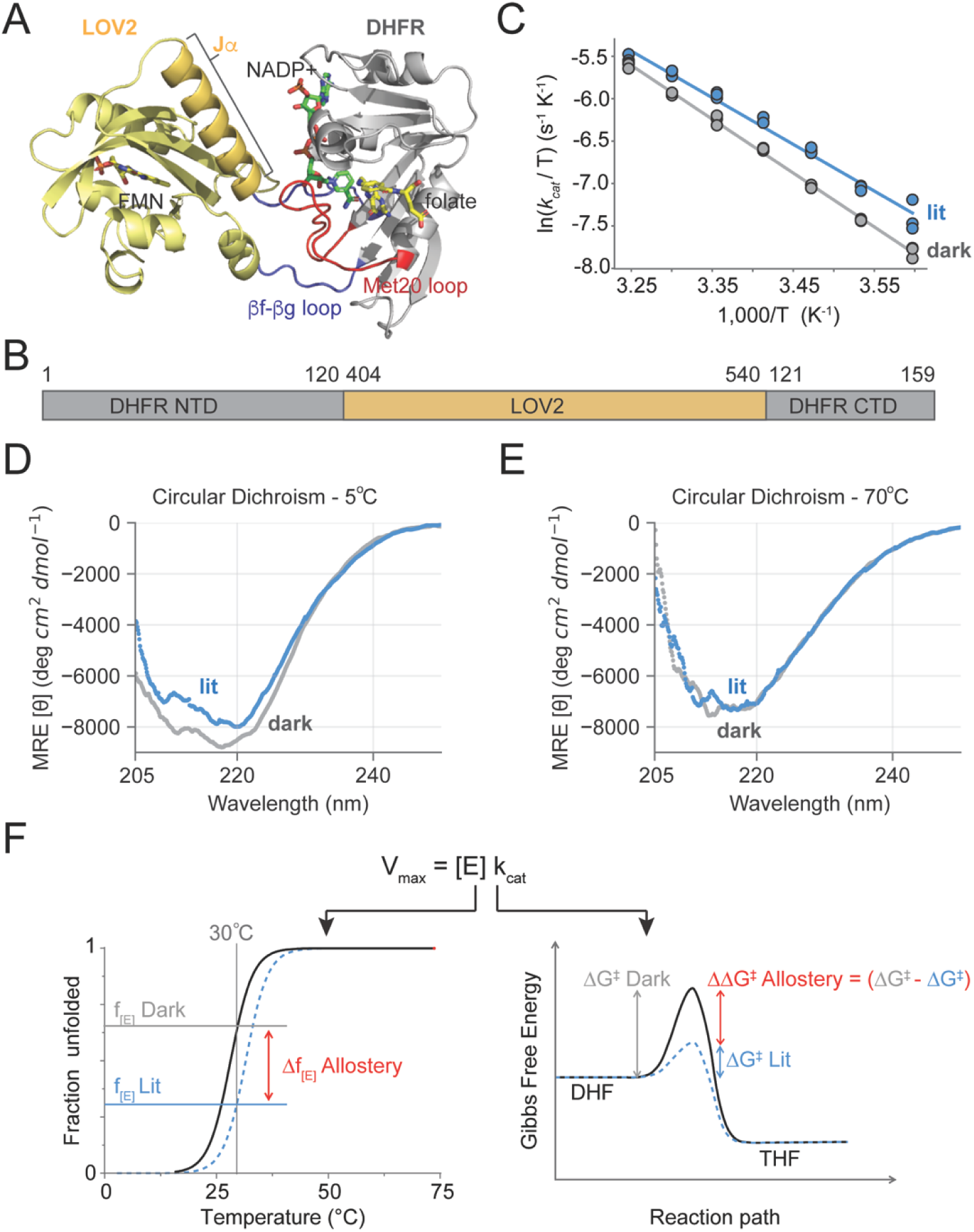
Light regulation of catalytic activity and structural disorder in the DL121 fusion. (A) AlphaFold model of the DL121 structure. The LOV2 domain (yellow cartoon) and DHFR domain (grey cartoon) are covalently linked. More specifically, LOV2 is inserted into the DHFR βF-βG loop (blue cartoon), which stabilizes the catalytically important Met20 loop (red cartoon) in the native enzyme. The FMN chromophore of LOV2 is indicated in yellow sticks, the NADP+ cofactor of DHFR in green sticks, and the folate substrate of DHFR in yellow sticks. (B) Domain structure and numbering of the DL121 fusion. (C) Temperature-dependent catalytic activity and allostery for the DL121 fusion. Each dot indicates a single experimental measurement of catalytic turnover (*k_cat_*) at 5°C intervals between 5-35°C. Blue points were measured in the light, grey points in the dark; three measurements were made at each temperature. Solid lines are the line-of-best fit for the Eyring equation across all 21 data points. (D) Circular dichroism measurements under lit (blue) and dark (grey) conditions for DL121 at 5°C. Solid lines represent an aggregate of triplicate scans. All measurements were in 100 mM NaF and 50 mM sodium phosphate at pH 6.5. (E) Circular dichroism measurements under lit (blue) and dark (grey) conditions for DL121 at 45°C. Solid lines represent an average across triplicate scans. All measurements were in 10 mM NaF and 50 mM sodium phosphate pH 6.5. (F) Schematic of thermodynamic forces shaping allosteric regulation. Temperature-dependent variation in allostery could reflect light-dependent changes in stability or the transition free energy.

## RESULTS

### DL121 shows temperature-dependent allostery

DHFR — the output domain for our engineered allosteric system — has become a model enzyme for studying the connection between conformational change and catalysis ^16–20^. DHFR catalyzes the stereospecific reduction of dihydrofolate to tetrahydrofolate using NADPH as a cofactor; tetrahydrofolate then serves as a carrier for one-carbon units in a variety of downstream reactions including the biosynthesis of thymidine, inosine (a purine precursor), glycine, and methionine ^21,22^. In the native *E. coli* enzyme, catalytic turnover proceeds through five chemical intermediates (Fig. S1A) ^23^. Prior structural studies have identified key loop motions associated with these intermediates that influence substrate binding, catalysis, and product release ^18,24^. A key feature of DHFR conformational change is the repositioning of the active-site adjacent Met20 loop from a catalytically competent “closed” state to an “occluded” state, a motion that is associated with cofactor exchange and subsequent product release (Fig.S1B,C). In the DL121 allosteric fusion, the LOV2 domain, comprised of residues 404-540 of *A. sativa* Phot1 kinase, are inserted within the βF-βG loop of DHFR (Fig. 1A, B). In the native DHFR enzyme, the βF-βG loop forms a pair of hydrogen bonds that stabilize the closed conformation of the Met20 loop (Fig. S1B). These hydrogen bonds are broken in the occluded conformation that follows hydride transfer (Fig. S1C).

We attempted crystallography for a variant of the DHFR/LOV2 fusion in which the LOV2 domain is mutated to not respond to light (DL121-C450S, sometimes referred to as a “dark-locked” mutation) ^5,25^. This construct resisted crystallization under both holo and apo conditions. However, an AlphaFold model provides an approximate sense of potential interdomain interactions (Fig. 1A). The βF-βG loop is substantially reorganized, and LOV2 is positioned adjacent to the NADP binding pocket of DHFR (Fig. 1A, Fig. S1D). Accordingly, the hydrogen bonds between the βF-βG and Met20 loops of DHFR are predicted to be disrupted (Fig. S1D). Consistent with this predicted structural rearrangement, we observed that DL121 is approximately 21-fold less active than the native enzyme: at 25°C the *k_cat_* for native DHFR and DL121 are 12.3 and 0.58 respectively. Under lit conditions (also at 25°C), DL121 undergoes an 34% activation in *k_cat_*, but no statistically significant change in *K_m_*. This is consistent with “V-type” allostery, in which the allosteric effector modulates the rate of catalysis rather than substrate binding ^15,26^. While product release is the rate limiting step for catalysis in the native DHFR, prior work suggests that the rate of hydride transfer is limiting for DL121 (Fig. S1A) ^5^. In agreement with this, we found that catalytic turnover is strongly pH dependent (Fig. S2).

To more completely understand the thermodynamic contributions to catalysis in the DL121 enzyme, we monitored both catalysis and conformational change as function of temperature. More specifically, we measured *k_cat_*in light and dark conditions as a function of temperature under saturating conditions of substrate and cofactor (Fig. 1C). We observed that catalytic activation in response to light was strongly temperature dependent, with allostery becoming more pronounced at low temperatures. This suggests that the allosteric effect is enthalpically driven, with an entropic penalty that increases with higher temperatures. Then, using circular dichroism (CD) spectroscopy, we monitored the light-dependent response of DL121 secondary structure. We used a fiber optic coupled blue LED to excite the DL121 fusion in a CD spectrophotometer and collected spectra across the near UV under both dark and lit conditions. At low temperatures (Fig. 1D), the loss of α-helical content in the 205-225 nm range can be seen when the protein is illuminated. This is consistent with the light-induced conformational changes of the wild type AsLOV2 domain where the Jα helix unfolds and is displaced from the PAS core^9^. However, this change is not observed at higher temperatures (Fig. 1E), presumably because the protein is now substantially more disordered. Thus, we observed that the conformational change in response to light is temperature sensitive much like the catalytic response to light. These data suggested two possible allosteric mechanisms for DL121: 1) that LOV2 regulates DHFR activity by simply impacting the fraction folded into a catalytically competent state, or 2) that LOV2 directly modulates energetics of the transition state. Given this, we next sought to answer whether the temperature-dependent variation in allostery reflects light-dependent stability changes or perturbations to the transition free energy (Fig. 1F).

### Light activation of DHFR is mediated by enthalpy-driven stabilization of the catalytic transition state

Next, we used Eyring analysis to calculate the transition state enthalpy and entropy under both lit and dark conditions from the *k_cat_* data in Fig 1C. To provide a more robust estimate of the error in our data, we used bootstrap resampling to compute 95% CI and estimates of standard error. We observed a modest enthalpic benefit of light activation (ΔΔ*H_Lit–Dark_*^‡^ = - 1665 cal/mol) that was partially countered by an entropic penalty (ΔΔ*S_Lit–Dark_*^‡^ = -1536 cal/mol at 30°C) (Fig. 2A,B, Fig. S3, Table S1).

**Figure 2.**
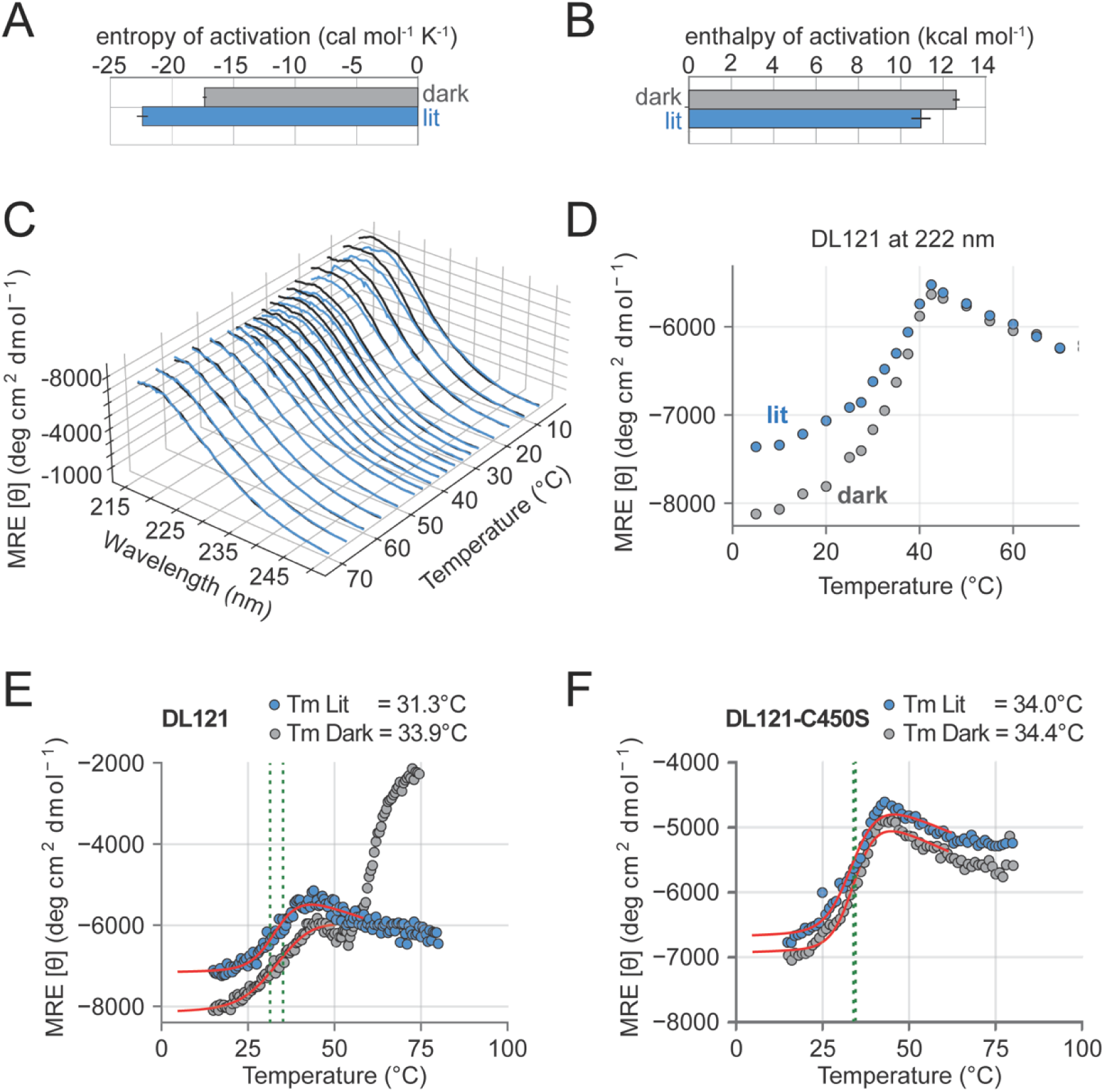
Thermodynamic effects of light on DL121 transition state thermodynamics and global stability. (A) Entropy of the DL121 catalytic transition state (Δ*S*^‡^) as inferred by Eyring analysis. All experimental measurements were made in technical triplicate under lit and dark conditions. Bar heights reflect the average estimated by bootstrap resampling; error bars are the corresponding standard deviation as estimated by bootstrap resampling. (B) Enthalpy of the DL121 catalytic transition state (Δ*H*^‡^) as inferred by Eyring analysis. All experimental measurements were made in technical triplicate under lit and dark conditions. Bar heights reflect the average estimated by bootstrap resampling; error bars are the corresponding standard deviation as estimated by bootstrap resampling. (C) Circular dichroism spectra for DL121 under lit (blue) and dark (grey) conditions as a function of temperature. (D) Molar ellipticity at 222 nm for DL121 under lit (blue) and dark (grey) conditions. Data are a subset of those in (C). (E) Thermal unfolding of DL121 as monitored by CD under lit (blue) and dark (grey) conditions. The red lines indicate the best fit three state unfolding model. Points indicate individual experimental measurements aggregated across three triplicate scans. (F) Thermal unfolding of DL121 as monitored by CD under lit (blue) and dark (grey) conditions. The red lines indicate the best fit three state unfolding model. Points indicate individual experimental measurements aggregated across three triplicate scans.

To understand the contribution of stability to allostery, we collected complete CD spectra under light and dark conditions across 22 temperatures from 5 to 90°C, with more dense sampling between 5-70°C (Fig. 2C, Fig. S4A). At low temperatures we observed a light-dependent change in the CD signal between 205-225 nm, consistent with light-induced loss of alpha helical content. Focusing on the signal at 222 nm alone shows that this light-inducible conformational change collapses near 40°C (Fig. 2D, Fig. S4B,C). However, this temperature is beyond the range of our Eyring analysis (which extended to 35°C), suggesting that the temperature-dependent loss in allosteric regulation of DHFR is not primarily driven by the loss of LOV2 alpha helical content and functionality.

To more quantitatively examine the thermodynamics of unfolding, we collected thermal melts by monitoring CD at 222 nm with increased temperature resolution. Under lit conditions, DL121 shows a two-state unfolding transition. In contrast, the dark state melt of DL121 exhibited a second transition just above 50°C at 222 nm (Fig. 2E). This transition was seemingly related to aggregation, as we observed a concomitant decrease in the HT voltage signal accompanied by a rapid increase in the CD signal (Fig. S5). There was also visible aggregation in the cuvette following the melt. We did not observe aggregation or this second CD transition in samples that were periodically illuminated (e.g. Fig 2C,D) or in DL121 C450S (Fig. 2F, Fig. S5). This suggested that transiently sampling the lit state and lack of a cysteine at position 450 are both protective against large scale (visible) aggregate formation. Interestingly, prior work from Sedlak and co-workers also reported an increase in 222 nm signal at 65°C (following an initial transition near 40C) for the AsLOV2 domain in isolation (at pH 7 and above) ^27^. This transition was more modest for the AsLOV2 C450A mutation at pH 7 (the condition closest to our own). Like our own study, Sedlak and colleagues observed precipitation at pH 5 and 6 (our own conditions were pH 6.5). Following the proposition that the second transition reflects an artifact of aggregation, we treated DL121 as following a two-state unfolding equilibrium below 50°C under both dark and lit conditions.

We computed a melting temperature (T_m_) for the lit and dark DL121 samples by fitting a two-state thermodynamic model that accounts for pre- and post-transition linear changes in ellipticity^28^. We could not compute a free energy of unfolding for either protein variant because the reaction was not reversible following heating to high temperatures (Fig. S4D). These data revealed that DL121 is marginally stable at 30°C under dark conditions (T_m_ = 33.9°C) and that light is modestly destabilizing (T_m_ = 31.3°C) (Fig. 2E, Supplemental Table 2). Moreover, the light-insensitive DL121 C450S variant does not show light-dependent stability (Fig. 2F, T_m_ = 34.0°C and 34.4°C in the light and dark respectively).

Taken together, our data indicate that the lit state is less thermally stable while being more catalytically active. Thus, the allosteric effect is not simply explained by light increasing the proportion of a more structured, globally folded and therefore active state — if anything the observed effect would seem to oppose allostery by decreasing the fraction folded at 30°C. We concluded that local unfolding of LOV2 is not regulating catalytic activity by coupling to the global folding equilibrium of the fusion but rather promotes an enthalpically favorable conformation of the transition state.

### Allostery-tuning mutations act through an enthalpy-entropy tradeoff

Next, we sought to understand how structurally distributed mutations in the DHFR domain tune allosteric regulation. In prior work, we used a growth-based functional screen to quantify the impact of all possible single mutations in the DHFR domain of DL121 on allostery. We identified 69 statistically significant allostery-tuning mutations; 13 of which disrupted or reduced the DL121 allosteric signal, while 56 were allostery-enhancing. The allostery-tuning mutations were structurally distributed throughout the DHFR domain and enriched on the solvent-exposed surface. For this study, we chose nine of these mutations for more detailed biochemical and thermodynamic characterization: A9N, M16A, M16P, G86K, D87A, R98M, D116M, H124Q, and D127W (Fig. 3A-C). These mutations were initially selected to be distributed on the DHFR structure (Fig 3B), however mapping the mutations to the AlphaFold structure suggests that many of them lie near a potential DHFR/LOV2 interface (Fig 3A). The allostery-disrupting mutation D87A is an obvious exception.

**Figure 3.**
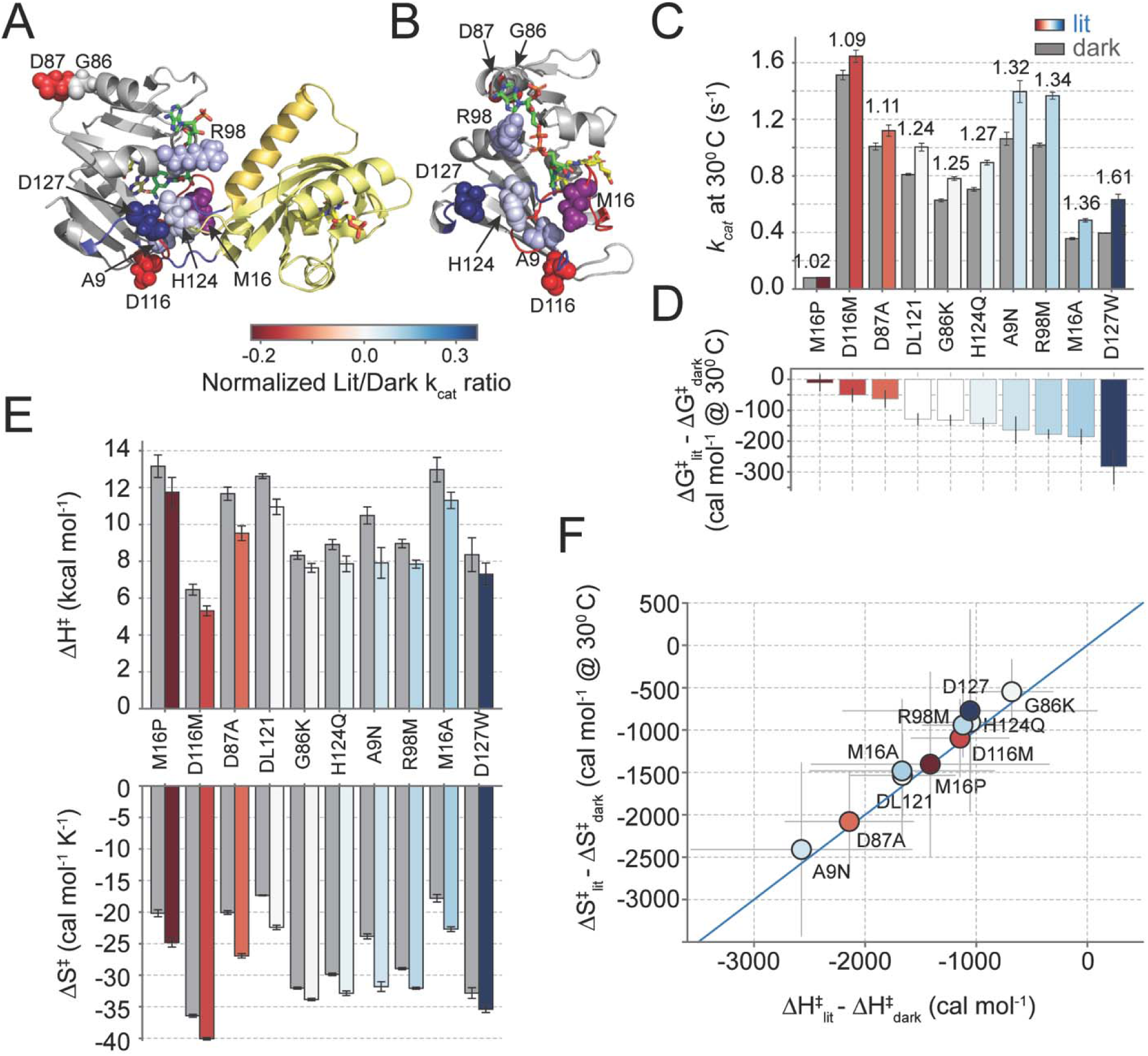
An enthalpy-entropy tradeoff in the catalytic effect of allostery tuning mutations. (A) The structural distribution of biochemically characterized allostery tuning mutations on an AlphaFold model of the DL121 fusion. The DHFR domain is indicated in grey cartoon, while LOV2 is in yellow cartoon. Mutated sites are shown as spheres and color coded by allosteric effect size (see legend below). Position 16 is colored purple, because mutations at this position had both allostery-enhancing (M16A) and disrupting (M16P) effects. (B) A second view of the allostery tuning mutations in the context of the DHFR domain only. Folate substrate (yellow) and NADP+ co-factor (green) are shown in sticks. Again, mutated sites are shown in spheres and color coded as in (A). (C) Catalytic turnover number for each mutant under lit (colored bars) and dark (grey bars) conditions. Bar height represents the average and error bars represent standard deviation as estimated across 5000 bootstrap re-samplings of triplicate experimental measurements across seven temperatures. Small numbers above each pair of bars indicate the allosteric fold change. The mutations are sorted by allosteric effect size. (D) The change in transition state free energy between the lit and dark states of the enzyme. Color coding follows from (A). Bar height represents the average and error bars represent standard deviation as estimated across 5000 bootstrap re-samplings of triplicate experimental measurements across seven temperatures. (E) Transition state enthalpy (top) and entropy (bottom) under lit (colored bars) and dark (grey bars) conditions as determined by Eyring analysis. Bar height represents the average and error bars represent standard deviation as estimated across 5000 bootstrap re-samplings of triplicate experimental measurements across seven temperatures. The mutations are sorted as in (C,D). (F) Correlation between the transition state enthalpy and entropy across DL121 and all nine mutations. The points are color coded as in (A) and reflect the average value. Error bars are shown as light grey lines and reflect propagated error from (E). The solid blue line indicates x=y, marking an exact entropy-enthalpy tradeoff.

Each DL121 mutant was expressed and purified, then *k_cat_*was measured with saturating substrate and cofactor under both lit and dark conditions. In comparison to our earlier work, we removed the histidine tag prior to catalytic characterization and added additional purification steps. Consequently, there are some differences in the catalytic efficiencies and allosteric effects measured here and in our 2021 manuscript (Fig. S6). Most notably, H124Q showed a more pronounced allosteric enhancement in our 2021 work in comparison to the present measurements. We attributed this variation to differences in purity but cannot exclude the possibility that the histidine tag itself contributed in some way to the earlier observed allosteric measurements.

The characterized mutations spanned a range of absolute effect sizes on catalysis and allosteric regulation (Fig. 3C). With the exception of M16P, all mutants retained some allosteric regulation. Two showed reduced regulation compared to DL121 (D116M, D87A), one was near-identical to DL121 (G86K), and the remaining five were allostery enhancing (A9N, M16A, R98M, H124Q, D127W). As in our prior work, there was no obvious correlation between the impact of mutation on catalysis and allosteric effect size ^15^. For example, M16P had a large effect on catalysis (relative to DL121) but minimal effect on regulation, while M16A had a more modest effect on catalysis while enhancing allosteric regulation.

We measured *k_cat_* across seven temperatures and performed Eyring analysis to quantify the enthalpy and entropy of the catalytic transition state for all 9 mutants (Fig 3D-F, Fig. S3, and Table S1). Like DL121, all mutants exhibited enthalpy-mediated allostery. That is, we observed an enthalpic benefit in the light that was partly offset by an entropic penalty (Fig. 3E). For M16P — the least allosteric variant — the offset in entropy and enthalpy was near-exact. The mutation had a large effect on both transition state enthalpy and entropy, but these effects opposed one another to limit the impact on allostery (Fig. 3D,E)

More generally, the allosteric effect size across all mutations was tuned by a strong enthalpy/entropy tradeoff. To visualize this, we constructed a correlation plot between the light-dependent change in transition state enthalpy and the light-dependent change in entropy (Fig. 3F). In this plot, mutations along the x=y line have near-exact entropy/enthalpy offsets and minimal allostery (like M16P). Allosteric effect size increases as mutations appear further above the x=y line (as exemplified by D127W). Because all mutants exhibited enthalpy-mediated allostery, the allosteric effect was temperature dependent for all variants, such that the entropic penalty dominated at higher temperatures and reduced the impact of light on activity. Nonetheless, individual mutations can tune this temperature dependence. The strongest temperature dependence was observed at the lower left in Fig. 3F, with less temperature dependence at the top right. For example, A9N has a small allostery enhancing effect that is maximized at lower temperatures, while D127W shows a larger allostery enhancing effect that is better preserved across temperatures (Fig. S3 B,J). Overall, the observed entropy-enthalpy tradeoff seems to limit the capacity of single mutants to generate large effects in allosteric regulation.

### The impact of allostery-tuning mutations on global stability is idiosyncratic

We additionally quantified the impact of all nine mutations on global thermal stability. (Fig. 4A, Fig. S7-S11, and Table S2). As for the initial DL121 fusion, we monitored ellipticity at 222 nm over a temperature range spanning at least 5-80°C and calculated a T_m_ of unfolding by fitting a two-state model. In general, the mutations had modest effects of a few degrees on T_m_, with M16P and A9N being among the most stabilizing and G86K being the most destabilizing. There was not a clear relationship between the impact of mutation on catalytic activity and stability. For example, G86K is substantially destabilizing but has little impact on catalytic turnover. While five variants (including DL121) were more stable in the dark than the light, five others inverted this relationship and were more stable in the light. Thus, the local unfolding of the Jα helix in the lit state did not always lead to a lower global stability of the DL121 chimera.

**Figure 4.**
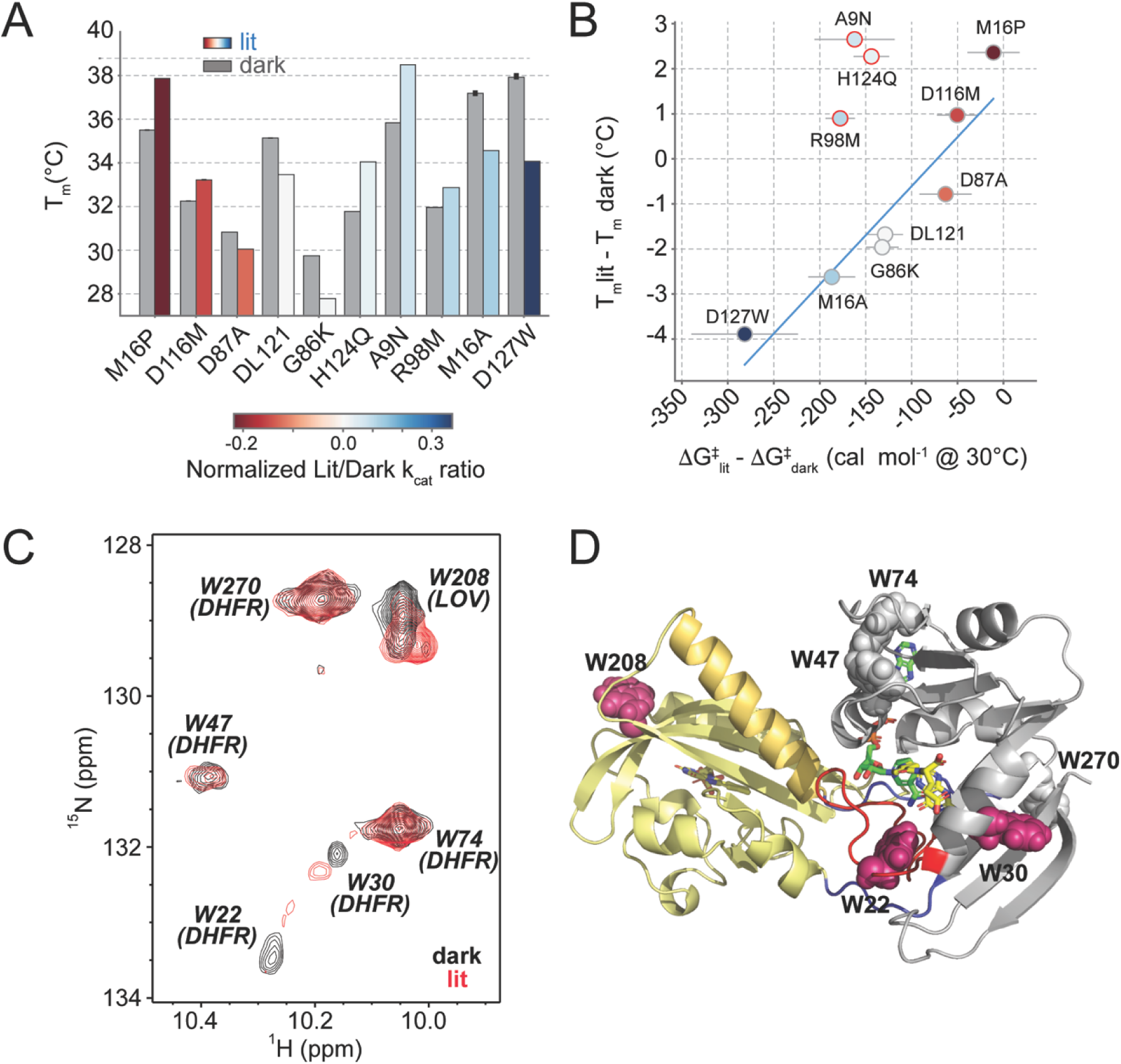
The relationship between global stability, local conformational flexibility, and transition state free energy. (A) Melting temperature under lit and dark conditions for selected allostery-tuning mutations (colored and grey bars respectively). Bar heights represent the best fit T_m_ for a two-state unfolding model. Bars are color coded according to the allosteric effect size of each variant. (B) The correlation between changes in thermal stability (y-axis) and the transition state free energy (x-axis). Each point corresponds to one mutation and is color coded as in (A). T_m_ values were measured in singlicate, transition state free energy values and error bars reflect the mean and standard deviation across 5000 bootstrap resamplings of triplicate experimental measurements across seven temperatures. The blue line is a line of best fit, and omitted the points outlined in red. (C) ^15^N/^1^H TROSY spectra of folate-loaded DL121 collected under dark (black) and light (red) conditions, utilizing 488 nm laser illumination to generate the photoexcited state *in situ*. (D) Structural location of the DL121 tryptophans. The structural model was generated by AlphaFold; DHFR domain is shown in grey and LOV2 in yellow. Tryptophans are shown in spheres, with the subset that show substantial light-dependent chemical shift changes associated with light are colored magenta.

To better examine the relationship between catalysis, global stability and allosteric tuning, we compared the impact of mutations on the light-dependent stability and transition state free energy of catalysis (Fig. 4B, Fig. S12). Light produced small effect sizes on stability and activity for all mutants (a few degrees for T_m_, and less than a *k_cal_* for the transition state free energy). Nonetheless, we observed that allostery-tuning mutations with the largest light-dependent effects on catalysis were negatively correlated to the impact on light-dependent global stability. That is, decreased stability in the light (relative to the dark) seemed to be associated with increased activity in the light (relative to the dark). This was true for the allostery-disrupting mutations (D87A, D116M, M16P) and the allostery-enhancing mutations (G86K, M16A, and D127W). However, a light-dependent change in T_m_ was not necessarily associated with a change in transition state free energy – three mutations were stabilized in the light relative to the dark and retained an allosteric effect size similar to DL121 (A9N, H124Q and R98M, highlighted with red circles in Fig. 4B). Thus, it was possible to alter the relative stabilities of the lit and dark states without tuning allostery. These three stability-tuning mutations with reduced allosteric effects lie near the putative DHFR-LOV2 interface in the AlphaFold structure, while the other mutations were seemingly more distal. It is possible that these mutations alter the DHFR-LOV2 interaction more directly without allosteric coupling to the Jα helix and βF-βG loop motions involved in catalysis. We never observed allosteric tuning without an accompanying stability effect, suggesting altered stability of the lit state may be necessary but not sufficient for a mutation to tune allosteric regulation.

### Solution NMR identifies light-induced chemical shift changes across both domains

To further probe the structural changes associated with LOV2 light activation, we collected 2D ^15^N/^1^H TROSY spectra of DL121 under dark and lit conditions in the presence of various substrates. Prior crystallographic work found that native DHFR occupies the occluded state when bound to folate, and the catalytically competent closed state when bound to folate and NADP+^24^. Importantly, we found that loading the complex with folate partly ordered the DL121 fusion, as indicated by improved homogeneity in peak width and intensity (Fig. S13A). Spectra in the presence of both folate and NADP+ somewhat reduced these improvements, consistent with a conformational change associated with loading both substrate and co-factor. Together, these data indicate that substrate binding may play a role in ordering the DHFR domain, which in turn promotes the capacity for allosteric signal transmission. We have observed similar phenomena in natural LOV-containing photosensors, including LOV-histidine kinases which have markedly improved LOV/effector coupling in the presence of ATP or other nucleotide analogs^29–31^.

Focusing on the folate-bound complex, we used a set of six tryptophan sidechain indole Nε1/Hε1 signals – one in LOV2, five in DHFR – in ^15^N/^1^H TROSY spectra as probes. We transferred these chemical shift assignments from BioMagResBank entries for *E. coli* DHFR and AsLOV2, using numbering scheme of DL121 (W208 = AsLOV2 W491; W270 = DHFR W133; all other tryptophans correspond to native DHFR numbering)^9,32^ and the observation that all six signals were only observed in the ligand-bound spectra. This included several peaks from sidechains which are proximal to folate (W22, W30, W47), consistent with ordering of the DHFR Met20 loop and nearby regions upon ligand addition. Of note, ligand binding within DHFR also influenced chemical shifts at the W208 sidechain within LOV2 (Fig. S13B), demonstrating bidirectional coupling between the two domains.

Light-induced chemical shift changes were observed throughout the ^15^N/^1^H TROSY spectra of DL121 in various liganded states, with specific changes clearest in presence of folate (Fig. S14). This included the LOV2-derived W208, as expected from prior studies of isolated LOV2^9^ (404-560) (Fig. 4C). Such light-induced chemical shift changes and broadening were not restricted solely to the LOV2 component of DL121, but also observed for DHFR-derived W22 and W30, consistent with transferred structural changes between the two proteins (Fig. 4C, 4D, S14B). Residue W22 is located in the DHFR Met20 loop, while W30 occurs near the folate binding pocket of the active site. No substantive changes were observed in the peaks for W47, W74, or W270. Together, these data suggest that light-induced structural changes in LOV2 propagate to the active site of DHFR but do not result in global changes in conformation or folding.

## DISCUSSION AND CONCLUSIONS

LOV2 domains have been used to introduce light regulation in multiple synthetic systems including a kinase ^6^, transcription factors ^33,34^, transcriptional repressor ^12^, and the Rac1 GTPase ^11,13^. Across these systems, the mechanism of regulation varies. Light has been used to alter localization, induce dimerization, “uncage” activity that was sterically occluded by the LOV2 domain, or reduce activity through light-induced local conformational disorder. Our results support yet another option for LOV2 regulation in engineered systems: that light-induced disorder can allosterically promote activity through increased flexibility. Blue light stimulation of DL121 resulted in a partial loss of secondary structure accompanied by a small reduction in thermal stability. This result excludes the possibility that light activation is simply mediated by an increase in global stability (and fraction folded) of DL121 within the cell. Instead, the structural changes in DL121 are associated with an energetically favorable decrease in catalytic transition state enthalpy that directly increases catalytic turnover.

Structural modeling suggests that unwinding of the LOV2 Jα helix may de-constrain key loop motions associated with DHFR catalysis. The LOV2 domain is inserted into the DHFR βF-βG loop, approximately 12 Å away from the active site. Interactions between the βF-βG and Met20 loops of the native DHFR enzyme are associated with a closed-to-occluded conformational transition of the Met20 loop that modulates NADP (cofactor) release ^18,24^. NMR relaxation dispersion experiments previously indicated Met20 closed-to-occluded conformational changes in controlling progression through the catalytic cycle ^35^. Moreover, atomic-resolution structural studies incorporating room temperature diffraction data and crystallography under electric field stimulation demonstrated a role for the Met20 loop in gating solvent access to the active site ^36^. Recent collaborative work involving some of the authors of this study employed replica-exchange molecular dynamics to examine how amino acid mutations at position 121 in DHFR modulate hydride transfer rate (*k_hyd_*) ^37^. The simulations revealed two Met20 loop conformations with subtle structural differences that altered substrate-cofactor positioning such that one conformation is catalytically competent while the other is not. Mutations at position 121 shifted the conformational equilibrium between these two states and consequently modulated hydride transfer. It is tempting to speculate that LOV2 insertion at position 121 similarly alters the Met20 loop dynamics in a light-dependent fashion. In any case, LOV2 fusion almost necessarily disrupts the native interactions between the βF-βG loop and the Met20 loop that shape conformational equilibria associated with catalysis. Though an atomically detailed structural explanation for the light-dependent increase in DHFR activity remains elusive, our NMR data confirmed bi-directional coupling between the two domains. Ligand binding within the DHFR active site modified chemical shifts in LOV2, and photoactivation of LOV2 modified chemical shifts in the active site of DHFR. Together these data indicate that a simple domain insertion (without further optimization of linker or interface) can introduce localized allosteric communication between an input effector and enzymatic active site.

Our data suggest that the dark state of LOV2 prevents the conformational fluctuations necessary to promote catalysis, while local unfolding in the lit state introduces the necessary flexibility to accommodate these changes. Consistent with this, we observed that allostery-disrupting mutations stabilize the lit state relative to the dark state. In contrast, allostery enhancing mutations with the largest effects further destabilize the lit state relative to the dark. Eyring analysis for a series of structurally distributed allostery-tuning mutations revealed that allosteric tuning is subject to a strong enthalpy-entropy tradeoff. For all mutants, the enthalpically favorable effects of light activation were opposed by an entropic penalty that limited the overall free energy impact of individual mutations on allostery. Depending on the balance of entropic and enthalpic contributions, the mutated DL121 variants showed modified temperature dependence of regulation. These results are strongly reminiscent of earlier work on enzyme thermal adaptation through surface mutations ^38^. For example, Saavedra et al. found that surface mutations that increased local disorder in Adenylate Kinase could be used to allosterically tune catalysis and ligand affinity ^39^. Moreover, molecular dynamics simulations of trypsin demonstrated that altering enzyme surface flexibility can tune the balance of enthalpic and entropic contributions to the catalytic transition state free energy, resulting in temperature adaptation ^40^. Our work analogously suggests that surface mutations can tune allosteric regulation (not just catalysis) by differentially modulating the balance of entropic and enthalpic contributions to the catalytic transition state under lit and dark conditions. We hypothesize that delocalized allosteric tuning is made possible by mutations that alter surface rigidity and interactions with the surrounding solvent, an idea that invites further testing with simulations, experiment, and analysis of extant thermally adapted sequences.

## MATERIALS AND METHODS

### Experimental model - *Escherichia coli* strains

XL1-Blue *E. coli* (genotype: *recA1 endA1 gyrA96 thi-1 hsdR17 supE44 relA1 lac* [F’ *proAB lacI*^q^*Z*Δ*M15* Tn*10*(Tet^r^)]) from Agilent Technologies were used for cloning, mutagenesis, and plasmid propagation. BL21(DE3) *E. coli* (genotype*: fhuA2 [lon] ompT gal (*λ *DE3) [dcm]* Δ*hsdS*. λ *DE3 =* λ *sBamHIo* Δ*EcoRI-B int::(lacI::PlacUV5::T7 gene1) i21* Δ*nin5*) from New England Biolabs were used for protein expression.

### DHFR Chimeric Expression Constructs

The *E. coli* DL121 fusion with N-terminal 8X His-tag in pHIS8-3 as described in McCormick et al. 2021 (addgene ID: 171953) was subcloned to improve the specificity and efficiency of tag removal. The new construct has a C-terminal 6X His-tag with TEV cleavage site and is expressed under control of a T7 promoter in the expression vector pET-24a. The C-terminal scar with a cut His-tag is GGGGSENLYFQ. Subcloning was accomplished by RF cloning (overlap extension PCR cloning); the DHFR LOV2 region was amplified from the original pHis8-3 construct by PCR with the primers ‘DL121-pet24a RF forward’ and ‘DL121-pet24a GS linker RF reverse’ as specified in Appendix 1^41^. Point mutants were engineered into the DHFR gene using QuikChange II site-directed mutagenesis (Agilent cat#200523) with primers specified in Appendix 1. All constructs were verified by Sanger DNA sequencing, and all DL121 fusions for purification in this work were expressed from this construct.

### Protein expression and purification

DL121 and mutant variants were expressed in BL21(DE3) *E. coli* grown at 30°C in Terrific Broth (12 g/L tryptone, 24 g/L yeast extract, 0.4% v/v glycerol, 17 mM KH_2_PO_4_, and 72 mM K_2_HPO_4_). Protein expression was induced with 0.5 mM IPTG when the cells reached an optical density of 0.7 at 600 nm, and cells were subsequently grown overnight at 18°C. The next morning, induced cells were pelleted (5000 RCF for 30 minutes at 4°C) and frozen in liquid nitrogen. Cell pellets were then thawed and lysed at 4°C by sonication in binding buffer added at a volume of 5 ml/g cell pellet (500 mM NaCl, 10 mM imidazole, 10 µg/ml leupeptin, 1 µg/ml pepstatin, 100 µM phenylmethylsulfonyl fluoride, 5 µg/ml DNase, 100 µg/ml lysozyme, 50 mM Tris-HCl, pH 7.0). Next the lysate was clarified by centrifugation (40,000 RCF for 30 minutes at 4°C) and the soluble fraction was incubated with Ni-NTA resin equilibrated in binding buffer (100 µL Ni-NTA slurry per gram of frozen cell pellet, Qiagen cat#4561) for 1 hour at 4°C. After washing the resin twice with one column volume of wash buffer (1M NaCl, 20mM imidazole, 50 mM Tris-HCl, pH 7.0) DL121 protein was eluted with two column volumes of elution buffer (1M NaCl, 400 mM imidazole, 100mM Tris-HCl, pH 7.0) at 4°C. Then 50 µg of His-tagged TEV protease per gram of cell pellet was added to the protein eluate and was dialyzed against 2 L of TEV cutting buffer (0.5 mM EDTA, 1 mM dithiothreitol, 50 mM Tris-HCl, pH 7.0) at 4°C overnight in a 10,000 MWCO Thermo protein Slide-A-Lyzer (Fisher Scientific cat#PI87730). The Slide-A-Lyzer was then moved into TEV binding buffer (1M NaCl, 5% glycerol, 50mM Tris pH 7) at 4°C for 4 hours. Following dialysis, the protein was again incubated with equilibrated Ni-NTA resin for 1 hour at 4°C to bind the His-tagged TEV and any uncut fraction of the protein (100µL Ni-NTA slurry per gram of frozen cell pellet, Qiagen cat#4561). The flowthrough containing the cut protein was then buffer exchanged into ion exchange A buffer (IEX-A, 10 mM β-mercaptoethanol, 50 mM Tris-HCl, pH 7.0) at 4°C using 3 rounds of dilution and concentration with an Amicon Ultra 10k M.W. cutoff concentrator (Sigma cat#UFC801024). The protein was then purified by ion exchange (1 mL HiTrap Q HP column, Cytiva cat#29051325) on a Bio-Rad NGC Quest 10 Plus Chromatography System. The column was run in linear gradient mode with increasing percentages of IEX-B buffer (10 mM β-mercaptoethanol, 1 M NaCl, 50 mM Tris-HCl, pH 7.0) over 10 column volumes up to 15% IEX-B buffer. At this point protein normally elutes. 15 column volumes of 15% IEX-B buffer are run in isocratic flow to allow full protein elution. The flow then returns to linear gradient mode up to 100% IEX-B buffer at a rate of 0.66 column volumes per percent increase of IEX-B buffer. The fractions containing protein were buffer exchanged into size exclusion FPLC buffer (SEFPLC buffer; 300 mM NaCl, 1% glycerol, 50 mM tris pH 7) and concentrated with a Amicon Ultra 10k M.W. cutoff concentrator (Sigma cat#UFC801024) to a volume of 2.5 mL. This concentrated sample was purified by size exclusion chromatography (HiLoad 16/600 Superdex 75 pg column, GE Life Sciences cat#28989333). Protein containing fractions of the eluate were tracked using the onboard multiwavelength UV/Vis detector set 280nm and 447nm. Purified protein was concentrated using Amicon Ulta 10k M.W. cutoff concentrator (Sigma cat#UFC801024) and flash frozen using liquid nitrogen. Purity of all samples was >95% by SDS-PAGE.

### Preparation of isotopically labeled DHFR

Isotopically labeled DHFR-LOV2 chimeric protein was expressed in BL21(DE3) *E. coli* grown in triple-labeled and supplemented M9 media (22mM Na_2_HPO_4_, 22mM KH_2_PO_4_, 8.5mM NaCl, 2 mM MgSO_4_, 18.5 mM ^15^NH_4_Cl (Cambridge Isotope Lab NLM-467-10), 16.5 mM ^13^C_6_-glucose (Cambridge Isotope Lab CLM 1396-26), 0.1 mM CaCl_2_, 0.036 mM FeSO_4_, 3µM Thiamine (CAS T1275-100G), 72 µM Kanamycin, using deuterium oxide as the solvent (Cambridge Isotope Lab DLM-4-1000, pH 7.4). Expression of deuterated protein was done as described in Li et al. 2022^42^. Briefly, 20 colonies were used to inoculate 15mL LB media in a 250mL flask. When the OD reached 0.6, 15ml of triple-labeled and supplemented M9 media was added. The cells continued to grow, and when the OD again reached 0.6, 30mL of triple-labeled and supplemented M9 media was added. Cell growth continued, and when OD reached 0.6, cells were pelleted by centrifugation at 5000 RCF for 10 minutes and resuspended in 100mL triple-labeled and supplemented M9 media which was allowed to grow overnight. The next morning 900mL triple-labeled and supplemented M9 media was added and to a 2.8L baffled shaker flask and shook at 30°C until the OD reached 0.7 whereupon protein expression was induced with 0.5 mM IPTG and cells were subsequently grown overnight at 18°C. The next morning induced cells were pelleted (5000 RCF for 30 minutes at 4°C) and frozen in liquid nitrogen. Protein purification proceed as described in the *Protein expression and purification* section.

### NMR data collection and analysis

All NMR spectra were acquired at 298 K on a 800.05 MHz Bruker Avance III HD spectrometer equipped with a 5 mm TCI CryoProbe, using samples of 320 µM U-^15^N labeled DL121 (with 1 mM folate and 1 mM NADP+, as indicated) in 50 mM sodium phosphate (pH 6.5), 100 mM NaCl, 1 mM EDTA, 1 mM DTT, protease inhibitor cocktail, and 10% D2O. ^15^N/^1^H WADE-TROSY spectra^43^ were collected on all these samples, integrating laser illumination (100 ms, 50 mW, 488 nm from Coherent Sapphire laser (Coherent, Santa Clara)) delivered via coaxial quartz fiber optic into the sample during the interscan recycle delay for lit state spectra. All data were processed using a combination of NMRPipe^44^, NMRViewJ^45^, and NMRFx Analyst^46^.

### Enzyme kinetic measurements

Purified DL121 protein was thawed at 4°C from -80°C frozen stocks. The protein was centrifuged at 21,130 RCF at 4°C for 10 minutes to remove any precipitate. The resulting supernatant was moved to a new tube with any pellet discarded. The concentration of the protein was quantitated by A_280_ using a DS-11+ spectrophotometer (DeNovix) with an extinction coefficient of 44,920 mM^-1^ cm^-1^ and MW of 34,889 Da. The *k_cat_* of DHFR was determined by monitoring the depletion of NADPH by absorbance at 340 nm (extinction coefficient = 13.2 mM^-1^ cm^-1^) under saturating concentrations of 25 µM DHF (Sigma-Aldrich cat#D7006) and 90 µM NADPH (Sigma-Aldrich cat#N7505). DHF substrate was prepared by suspension in MTEN buffer pH 7.00 with 0.35% β-mercaptoethanol and quantitated by A_282_ with an extinction coefficient of 28 mM^-1^ cm^-1^. All reactions were run in MTEN buffer pH 7.00 (50 mM 2-(N-morpholino)ethanesulfonic acid, 25 mM Tris, 25 mM ethanolamine, 100 mM NaCl), with 5 mM dithiothreitol. NADPH oxidation was monitored in 1-mL quartz cuvettes with a path length of 1 cm in a Lambda 650 UV/VIS spectrometer (Perkin Elmer) with attached water Peltier system set to the appropriate temperature. Eyring analysis sampled temperatures of 5, 10, 15, 20, 25, 30, 35 and 40°C. Protein samples and DHF solution were kept on ice prior to assay, and moved from an ice water slurry to the water bath/Peltier system 5 minutes before the reaction was begun. All samples were illuminated by a full spectrum 125-watt 6400K compact fluorescent bulb (Hydrofarm Inc. cat#FLC125D) suspended 12 inches above the water bath. Lit samples were in transparent 1.5-ml tubes while dark samples were kept in opaque LiteSafe Black Microtubes (Argos cat#T7100BK). Reaction initiation and transfer of the sample from the microtube to the temperature acclimated cuvette in the spectrometer was done rapidly by manual pipetting. All lighting except for a darkroom lamp was transiently extinguished during this step for the dark samples. The 40°C measurements were not included in subsequent data analysis because the T_m_ of the chimeric protein and mutants was less than 40°C and the relationship between catalytic activity and temperature was no longer linear, suggesting denaturation. The first 15 seconds of the reaction was used to calculate the initial reaction velocity by linear regression. The data across all temperatures was interpreted with Eyring analysis to estimate the enthalpy and entropy of activation:

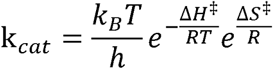

In this equation, *R* is the gas constant, *k_B_* is the Boltzmann constant, *h* is the Planck constant, Δ*H*^‡^ is enthalpy of activation, Δ*S*^‡^ is entropy of activation, and T is the temperature in Kelvin ^47^. The statistics of error for the activation energy were calculated using bootstrap sampling with replacement from the 21 k*_cat_* measurements (3 replicates for each of 7 temperatures) with n=5000 iterations.

### pH-dependent enzyme kinetics assay

DL121 *k_cat_* measurements were made as described above, but in MTEN buffer at pH of 5, 6, 7, 7.5, 8, 8.5, 9, and 9.5. The starting pH of a stock solution of MTEN was 6.00 and adjusted appropriately with HCl and NaOH. All samples were assayed at 17°C with 25 µM DHF, 100 nM protein, and 90 µM NADPH. The linear portion of the velocity curve for the first 15 seconds of the reaction was used to calculate the initial reaction velocity by linear regression.

### Spectrophotometry of the LOV2 chromophore

The lit- and dark-state spectra of the LOV2 chromophore were determined with a Lambda 650 UV/VIS spectrometer (Perkin Elmer) in scan mode at 350-550 nm using paired quartz cuvettes (Sigma cat#Z600350). Purified protein was diluted to 20 μM (unless otherwise noted) in MTEN buffer pH 7.00. The lit samples were illuminated for at least 2 minutes by full spectrum 125-watt 6400K compact fluorescent bulb (Hydrofarm Inc. cat#FLC125D). Relaxation of the lit-state chromophore was observed using the same samples and conditions as for the spectral measurement but with the instrument in continuous measurement mode at 447 nm. The relaxation constant was estimated from a one-phase association model using non-linear regression in GraphPad Prism 10.

### Circular Dichroism spectroscopy

CD spectroscopy was done on a Jasco J-815 CD Spectrometer. Protein samples were thawed and spun down at 21,130 RCF at 4°C for 10 minutes and the supernatant was moved to a new tube with any pellet being discarded. The concentration of the protein was quantitated by A_280_ using a DS-11+ spectrophotometer (DeNovix) with an extinction coefficient of 44920 mM^-1^ cm^-1^ and MW of 34,889 Da. The protein was then buffer exchanged into 10 mM NaF and 50 mM sodium phosphate pH 6.5 and concentrated with an Amicon Ulta 10k M.W. cutoff concentrator (Sigma cat#UFC801024). Samples were concentrated to 1 mg/ml, 300 μL of which was loaded into 1 mm path length quartz cuvettes. Samples were illuminated while in the instrument using a 455 nm, 1000 mA, Fiber-Coupled LED (ThorLabs #M455F3) fed through the injection port and secured into the top of the cuvette using a 200-uL pipette tip. The 455 nm LED was powered by a high-power 1-Channel LED driver (ThorLabs #DC2200) set to 900 mA current limit, brightness 100%, and a pulse of 250 ms with a total time of 10 s between each pulse. For lit samples, the LED controller was set to on, for dark samples the led controller was set to off. Unless otherwise noted, for all thermal protein melts the spectral bandwidth was set to a gap width reading of 1 nm at 222 nm and 222.1 nm, with 0.5°C/m ramp, 1 minute wait time, and 1 s DIT time. CD measurements that coincide with light pulses were computationally identified by a steep drop in the HT signal and were removed from the dataset. For spectral scans, an average of three scans were taken to prevent data gaps from light pulses. For the melt assays, CD values of dual measurements at 222 nm and 222.1 nm were averaged for the same reason. To calculate the T_m_ from the thermal melt data, we fit a two-state transition model as a function of temperature. The model assumes a monomer transitioning between folded and unfolded forms with corrections for pre- and post-transition linear changes in ellipticity ^28^:

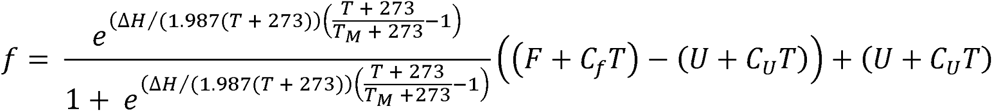

In this equation, *f* is fit to the CD measurement *θ*. Δ*H* is the enthalpy of folding, *T* is temperature in °C, T*_M_* is the temperature at which half the protein is melted in °C, *F* is mean residue ellipticity at 100% folded, *C_f_* is the linear correction folded function of temperature, *U* is mean residue ellipticity at 100% unfolded, and *C_u_* is the linear correction unfolded function of temperature.

As most of the protein samples precipitated over the course of a dark melts (but not in the light) we sought to exclude erroneous CD measurements caused by protein precipitation. As a decrease in photo-multiplier voltage value (HT) with increasing temperature is indicative of protein precipitation, we used the data from the HT signal to inform the bounds of the fit ^48,49^. This was confirmed by a visual inspection into the cell following each experiment. The starting point of the fit was the lowest experimentallymeasured temperature. The endpoint of the fit was defined as the halfway point between the maximum CD value and the maximum HT value. The code implementation of the light detection and unfolding model curve fitting is available on GitHub (https://github.com/reynoldsk/DHFRLOV121_thermo).

### Prediction of DL121 Structure with AlphaFold2

The structure of unmutated DL121 was predicted using the Astrocyte AlphaFold Workflow 0.0.2 developed by UT Southwestern Medical Center BioHPC. The workflow was run with reduced databases and a maximum template date of September 30, 2021. For MSA generation, 80,799 hidden Markov models (HHMs) were searched, resulting in a multiple sequence alignment comprising 421 sequences, with an effective sequence count of 5.39486. Five similar structural models were produced and ranked. The highest ranked model was selected. No additional post-processing steps were applied to the selected model.

**Table 1.**
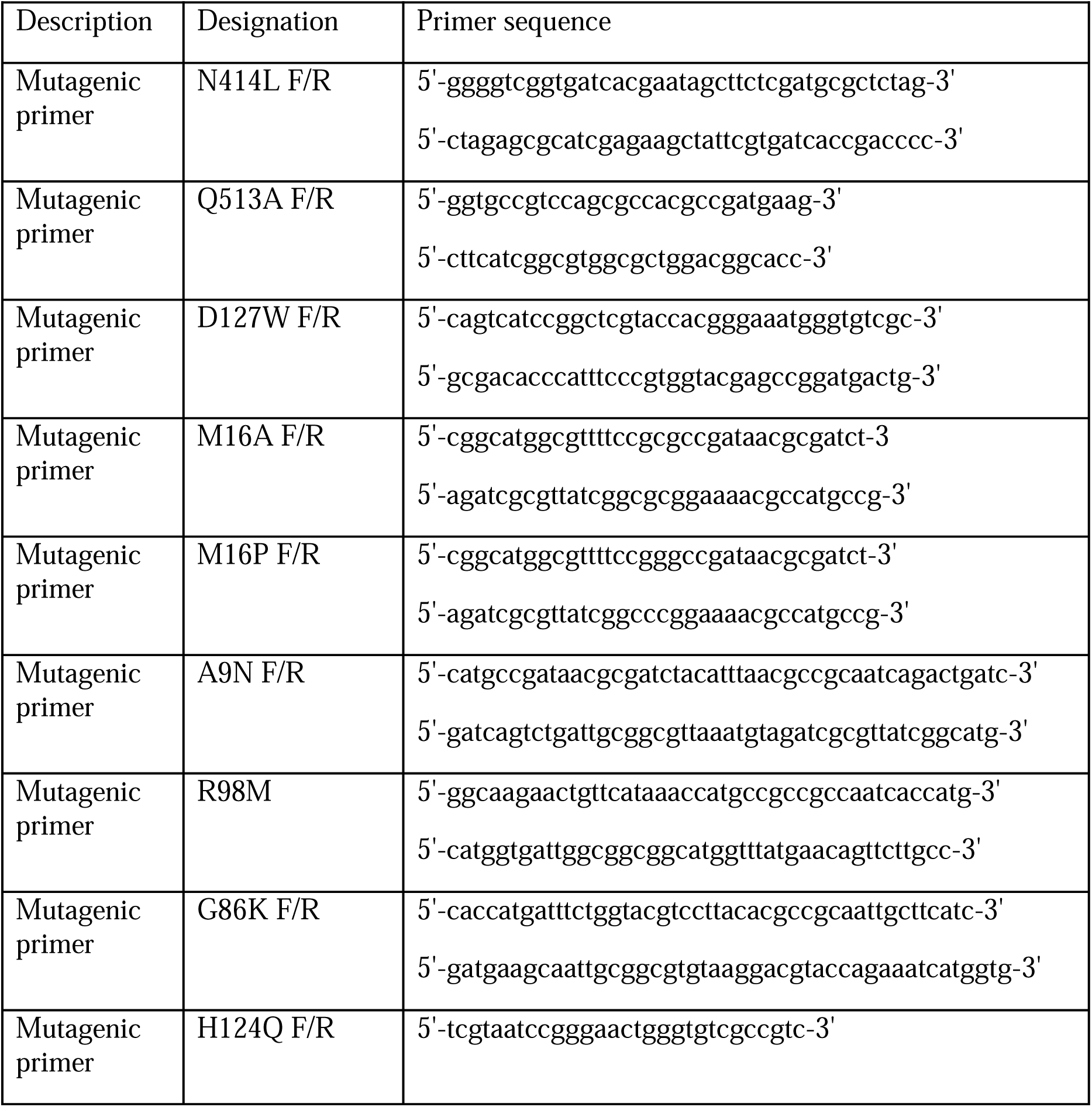
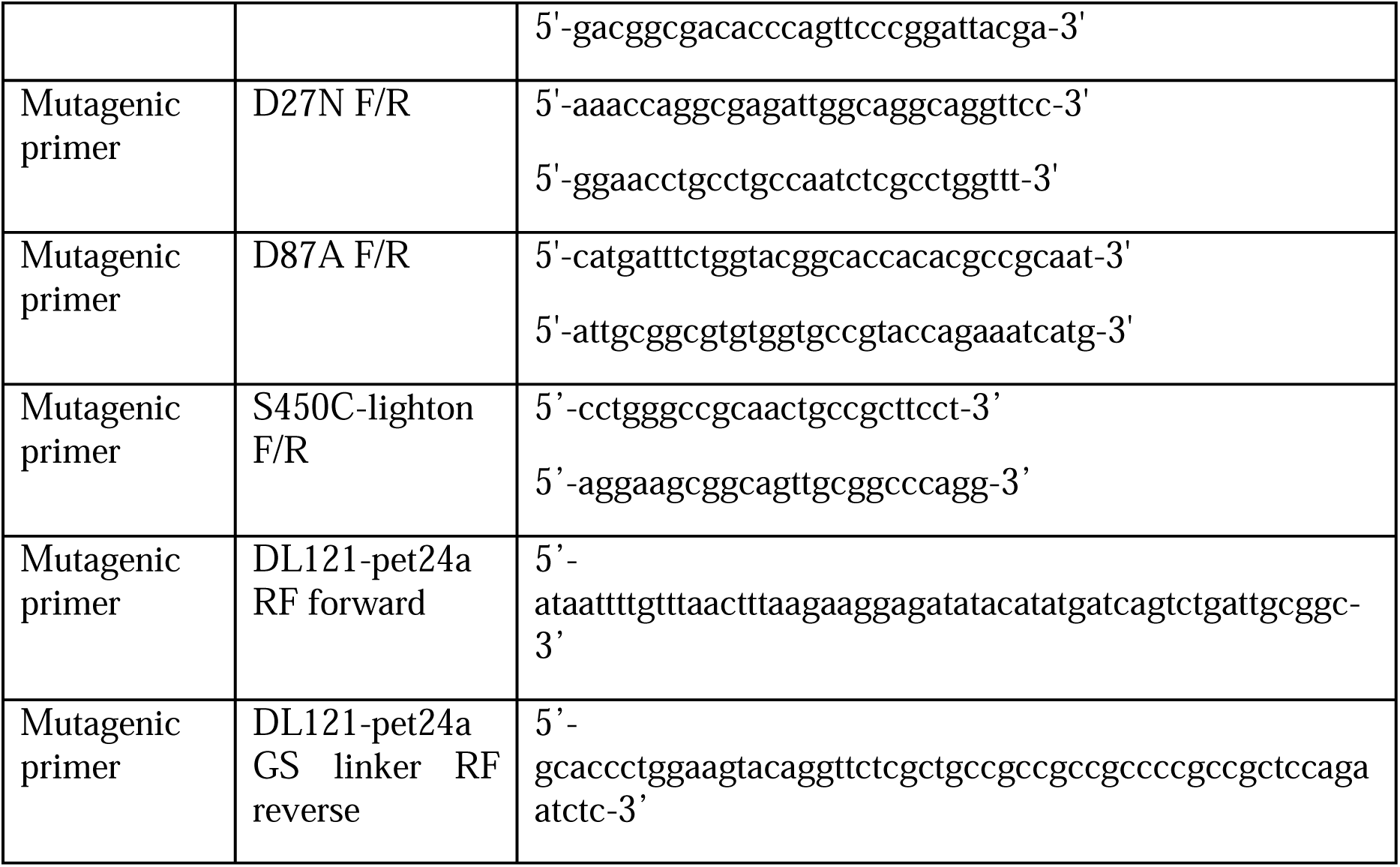
Primers used.

## Supporting information

Additional experimental details and data, including complete Eyring analysis, extended CD spectra, and thermal melts for all point mutants. These are presented as supplemental figures S1-S14 as well as supplementary tables S1 and S2 in SI.docx

## Funding Sources

This work was supported by NSF Grant #1942354 to KAR, NIH Grant R35 GM156296 to KHG, and NIH T32GM131963 to Jerry Dinan.

## Author Contributions

The manuscript was written through contributions of all authors. All authors have given approval to the final version of the manuscript.

## ACKNOWLEDGEMENTS

The authors thank Rama Ranganathan for early feedback and thoughts on this project. We thank Christine Ingle for her assistance with DHFR purification and kinetics protocols, Denize Favaro for assistance in collecting NMR spectra, Kendra Frederick for her feedback on our NMR work, Carla Madrid and Qiong Wu for help with identifying the appropriate NMR conditions, Chad Brautigam and Shih-Chia Tso for help with understanding our CD measurements, and fellow members of the Reynolds lab for feedback throughout the development of this work.

## DATA AVAILABILITY

The data and analysis underlying this study are openly available in github at: https://github.com/reynoldsk/DHFRLOV121_thermo

## Notes

https://github.com/reynoldsk/DHFRLOV121_thermo

### Competing Interest Statement

The authors have declared no competing interest.

### Summary of Updates

In this version, we have added new solution NMR data collected in collaboration with Dr. Kevin Gardner. These data are presented in new main text and supplementary figures (Figure 4, Figure S13) and a new results section entitled "Solution NMR identified light-induced chemical shift changes across both domains". We have also updated the methods section accordingly.

