## Supplemental Information for "Local disorder is associated with enhanced catalysis in a nascent photoswitch"

### **Supplemental Information for:**
### Local disorder is associated with enhanced catalysis in a nascent photoswitch

**Contents:**

**I. Supplementary Figures**


**Figure S1:** Overview of the DHFR catalytic cycle and associated conformational changes

**Figure S2**: The pH dependence of catalytic turnover

**Figure S3:** Eyring analysis for DL121 and select allostery tuning mutations

**Figure S4**: Extended DL121 CD spectra - additional temperatures and reversibility of unfolding

**Figure S5:** Thermal melts and associated HT voltage signal for DL121 and DL121 C450S

**Figure S6**: Comparison of enzyme catalytic turnover ($k_{cat}$) values as measured in this study with prior functional screening and biochemical characterization.

**Figure S7**: Thermal melts and associated HT voltage signal for DL121 A9N and DL121 M16A

**Figure S8**: Thermal melts and associated HT voltage signal for DL121 M16P and DL121 G86K

**Figure S9**: Thermal melts and associated HT voltage signal for DL121 D87A and DL121 R98M

**Figure S10**: Thermal melts and associated HT voltage signal for DL121 D116M and DL121 H124Q

**Figure S11**: Thermal melts and associated HT voltage signal for DL121 D127W

**Figure S12**: Relationship between thermal stability and transition state enthalpy and entropy

**Figure S13**: DL121 NMR chemical shift changes associated with folate binding.

**Figure S14:** Light-induced NMR chemical shift changes in DL121.


**II. Supplementary Tables**


**Table S1:** Transition state thermodynamic quantities and catalytic turnover measurements for DL121 and select allostery tuning mutations

**Table S2:** CD-derived thermal stability parameters for DL121 and select allostery tuning mutations

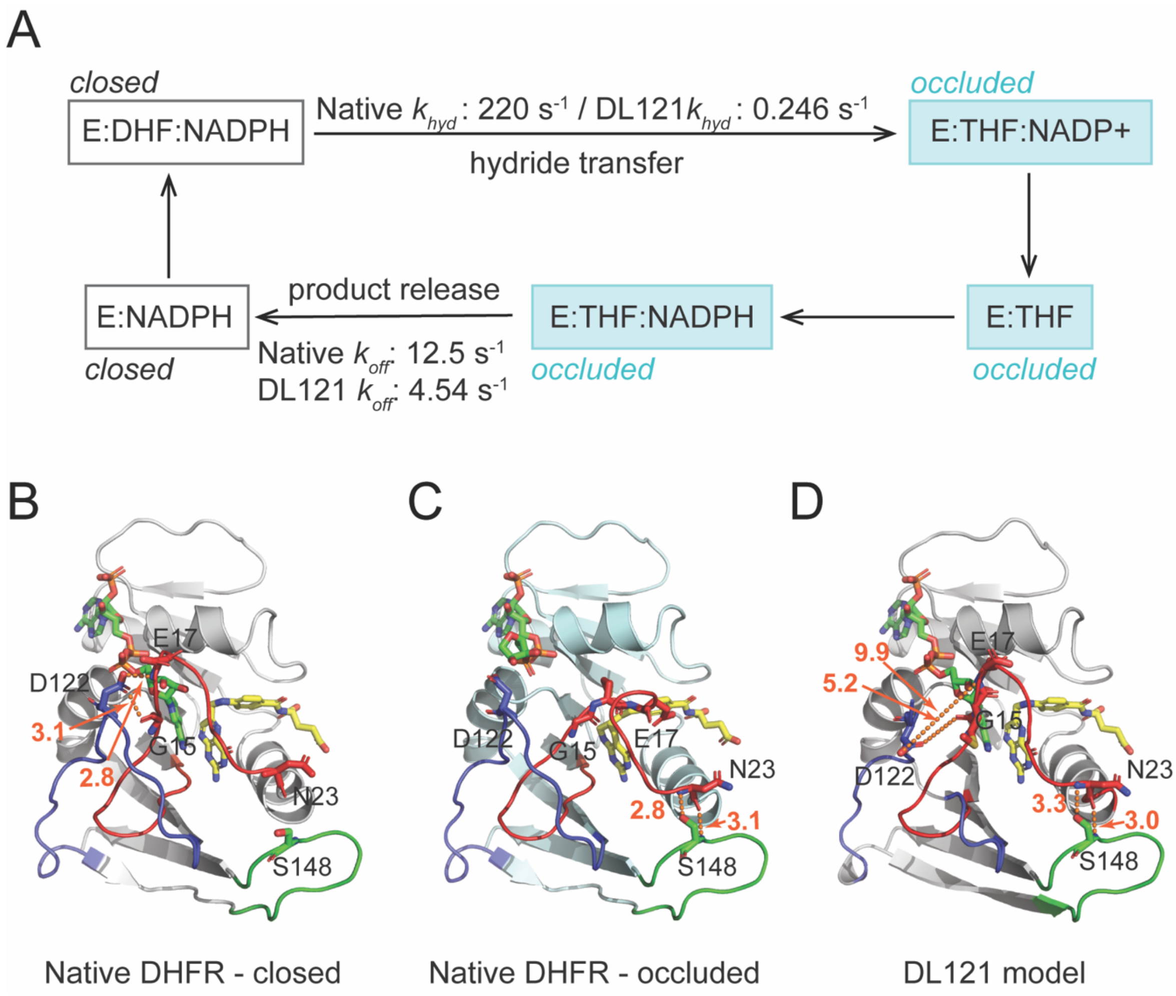


**Figure S1. Overview of the DHFR catalytic cycle and associated conformational changes**.

**A)** The DHFR catalytic cycle. Each square indicates a catalytic intermediate state, “E” stands for enzyme. Catalytic intermediates that adopt the occluded conformation are colored blue, while intermediates that adopt the closed conformation are colored white.

**B)** Key hydrogen bonding interactions in the closed conformation of native DHFR (PDB ID: 1RX2). The DHFR backbone is shown in cartoon, the DHF substrate in yellow sticks, and the NADP+ cofactor in green sticks. The βf-βg loop is shown in blue, the Met20 loop is colored red, and the G-H loop is green. Hydrogen bonds between the backbone amide of D122 and the backbone CO of Gly 15, as well as the O$\delta$ of D122 and backbone NH of Gly 17 are shown in orange dashed lines with distance in Angstroms indicated.

**C)** Key hydrogen bonding interactions in the occluded conformation of native DHFR (PDB ID: 1RX4). Hydrogen bonds between the backbone CO and NH of Asn23 and Ser148 (in the G-H loop) are shown in orange dashed lines with the distance in Angstroms indicated. Color coding is identical to (**B**).

**D)** Loop conformations in the Alpha-fold model of DL121. The LOV2 domain (between residues 120-121 of DHFR) is not shown to allow better visualization of the loops. The distances between the atoms that hydrogen bond in the closed and occluded conformations of the native enzyme are highlighted in orange dashed lines and labeled with the distance in Angstroms.

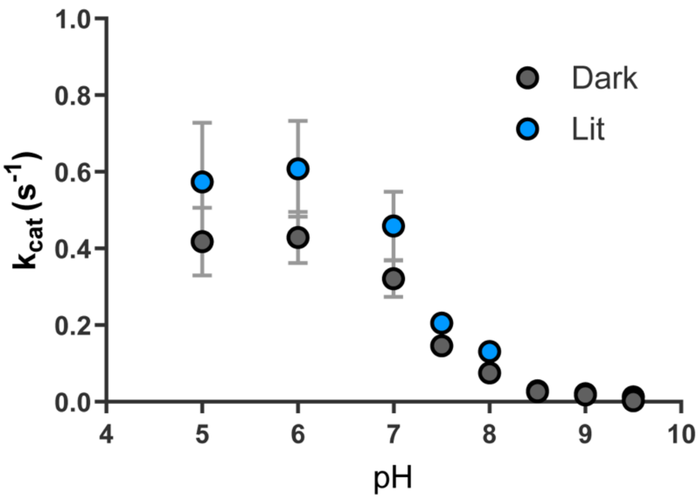


**Figure S2**: **The pH dependence of catalytic turnover (*$k_{cat}$*)**. Each point indicates the average of triplicate measurements in the light (blue) or dark (grey). Error bars are the standard deviation.

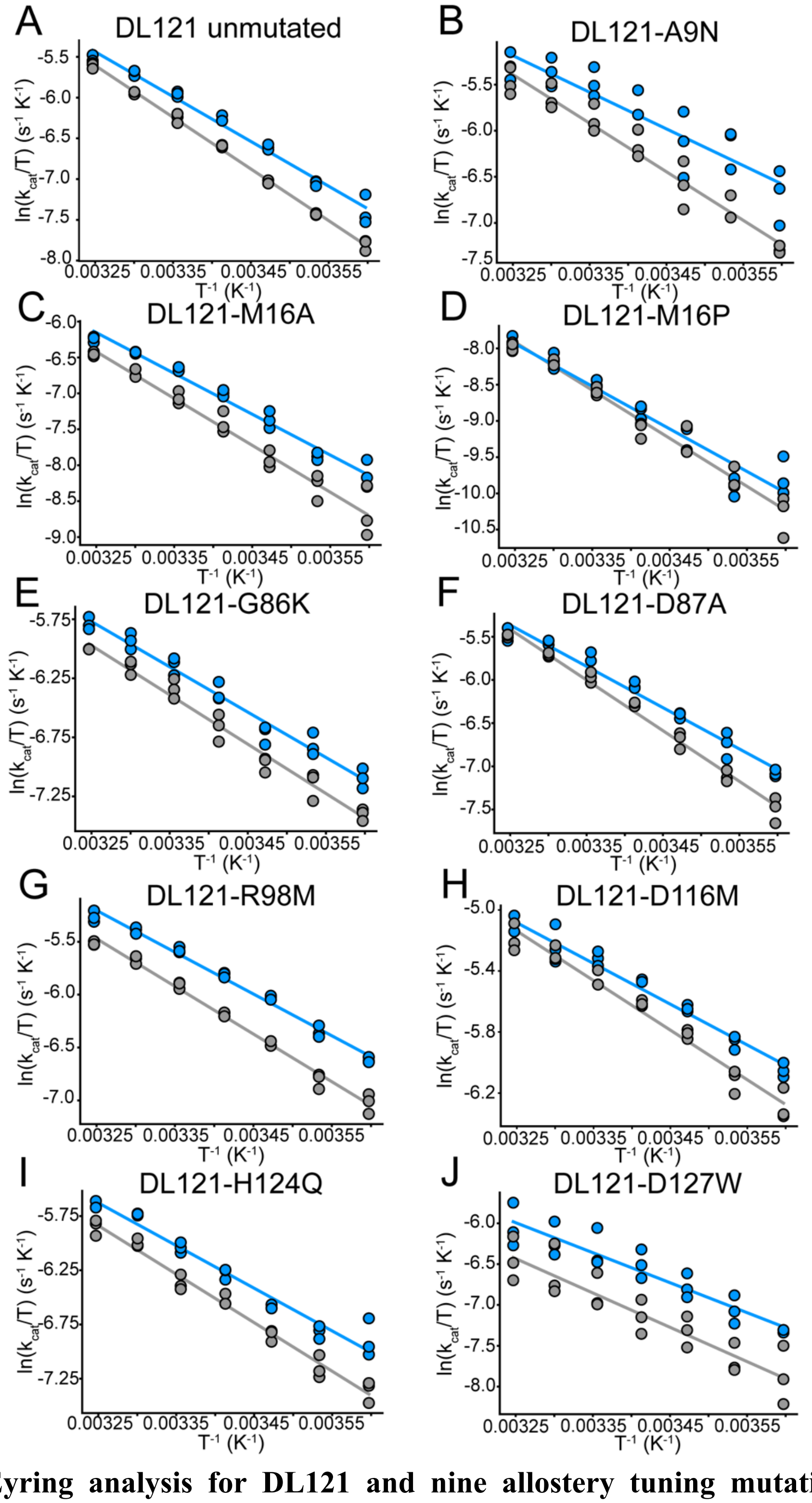


**Figure S3: Eyring analysis for DL121 and nine allostery tuning mutations**. Each point represents a single experimental measurement, catalytic turnover was measured in triplicate at seven temperatures in both the light (blue) and dark (grey). The lines represent the lines of best fit for the Eyring equation.

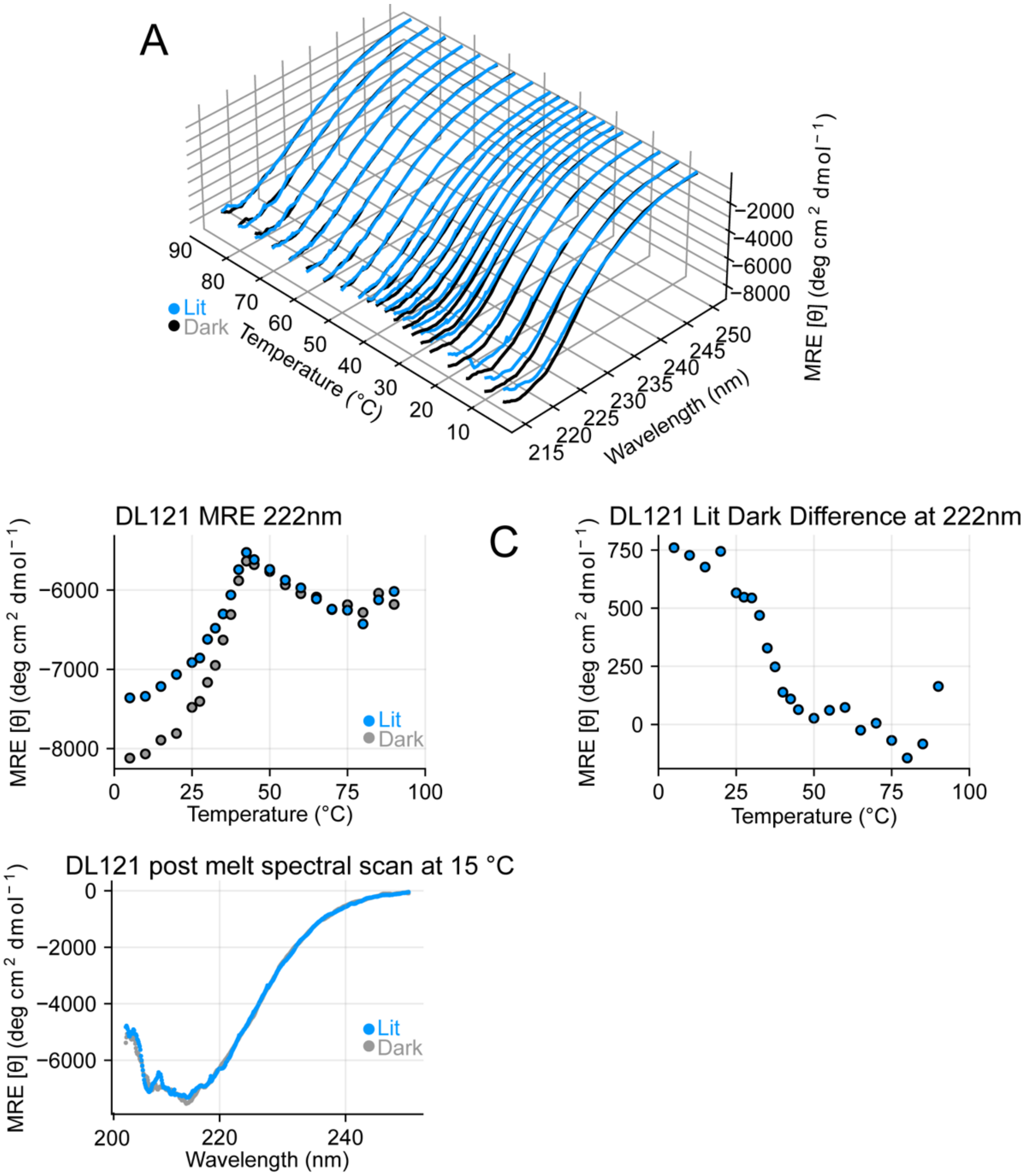


**Figure S4**: **Extended DL121 CD spectra - additional temperatures and reversibility of unfolding**.

**A)** Circular dichroism spectra for DL121 under lit (blue) and dark (grey) conditions as a function of temperature. Measurements were made at 21 temperatures from 5-90°C. These data are a superset of what is shown in Figure 2C.

**B)** Molar ellipticity at 222 nm for DL121 under lit (blue) and dark (grey) conditions. Data are a subset of those in (**A**).

**C)** The temperature dependent difference between molar ellipticity in the light and dark for DL121. Each point represents the difference of the data in (**B**).

**D)** Circular dichroism spectra for DL121 in the light and dark at 15°C following thermal unfolding (heating to 95°C). The spectra are consistent with an irreversible unfolding transition.

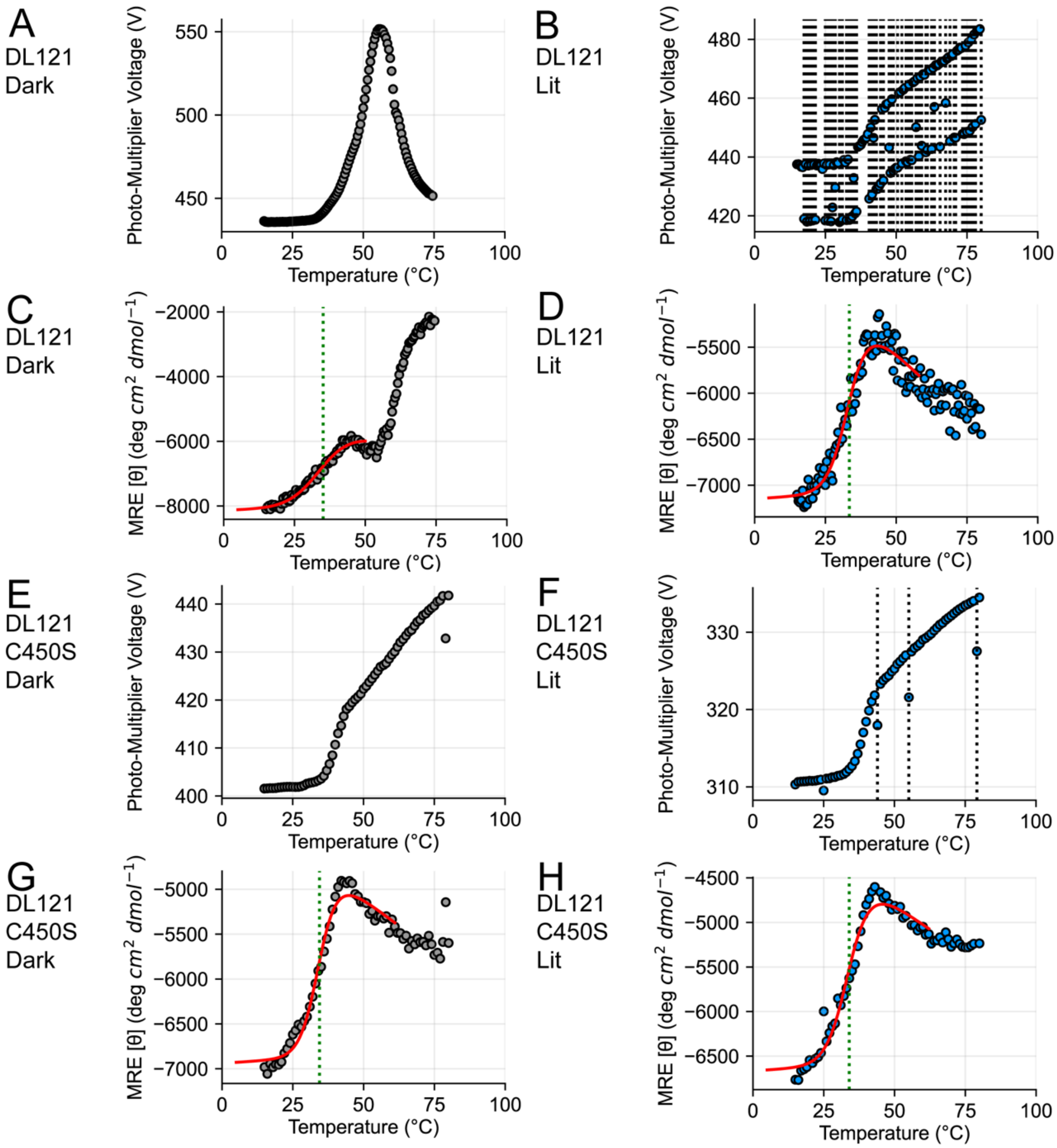


**Figure S5: Thermal melts and associated HT voltage signal for DL121 and DL121 C450S.**

**A)** Temperature dependence of the HT voltage signal for DL121 under dark conditions.

**B)** Temperature dependence of the HT voltage signal for DL121 under lit conditions. Dashed lines indicate blips in the signal accompanying light exposure, these were automatically detected from the HT signal and the accompanying points were removed from the molar ellipticity measurements in (D).

**C)** Temperature dependence of the molar ellipticity of DL121, measured in the dark. A dashed green line indicates the $T_m$, and the smooth red line is the best fit for a two state unfolding model that includes linear changes in ellipticity both pre- and post-transition.

**D)** Temperature dependence of the molar ellipticity of DL121, measured in the light. A dashed green line indicates the $T_m$, and the smooth red line is the best fit for a two state unfolding model that includes linear changes in ellipticity both pre- and post-transition.

**E-H**) Identical to A-D, but for DL121-C450S.

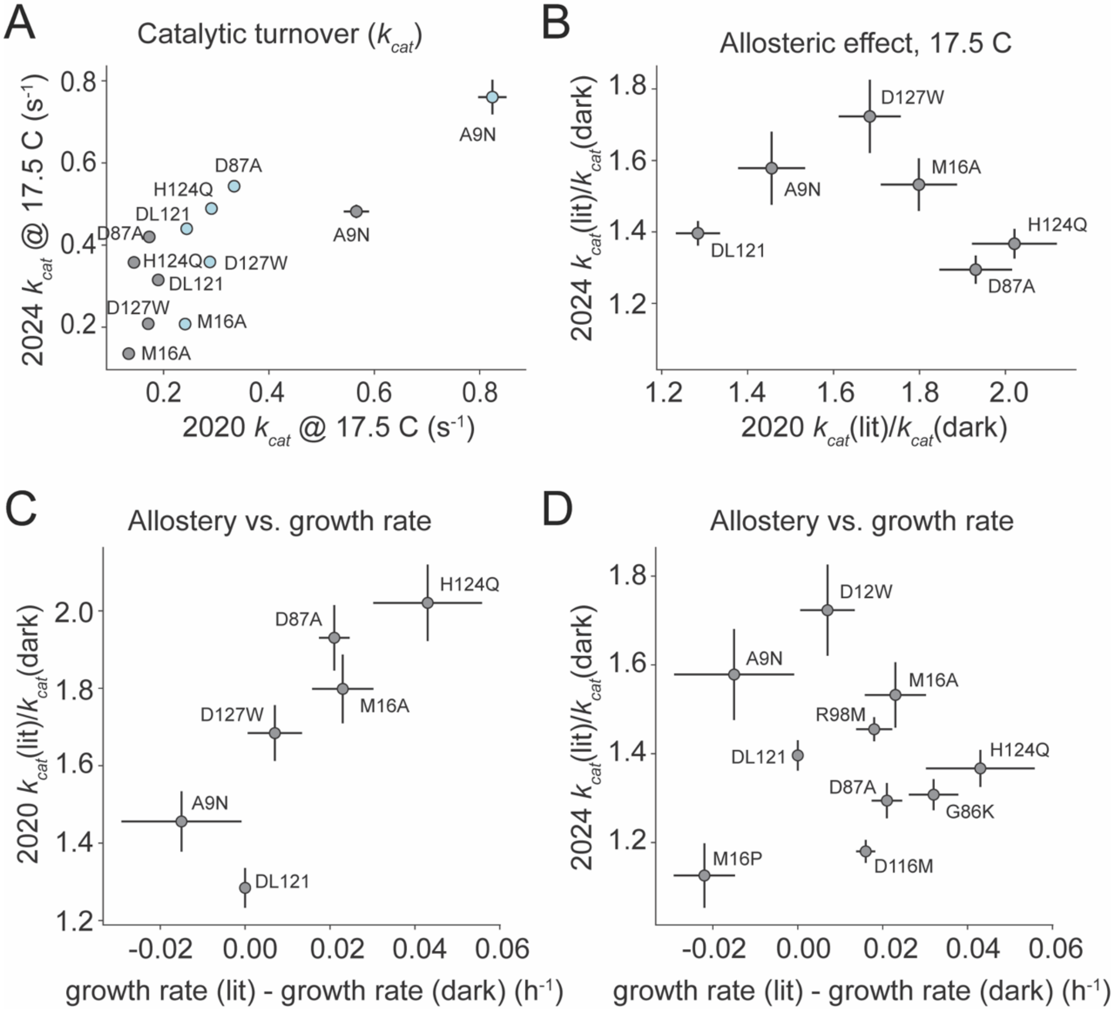


**Figure S6**: **Comparison of enzyme catalytic turnover (*$k_{cat}$*) values as measured in this study with prior functional screening and biochemical characterization**.

**A)** Correlation between $k_{cat}$ as measured in McCormick et al 2021 and the current work (McCormick et al 2024). Individual points represent the average of three experimental measurements made under both light (blue) and dark (grey) conditions. Error bars indicate the standard deviation across triplicates. Mutant names are indicated adjacent to each point.

**B)** Correlation in allosteric effect (the ratio of $k_{cat}$ in the light to $k_{cat}$ in the dark) as measured in 2024 and 2021. Each point represents the average of triplicate measurements, the error bars reflect propagated error from (**A**).

**C)** The relationship between the allosteric effect and the growth rate difference under lit and dark conditions in a functional screening assay, as reported in McCormick et al 2021.

**D)** The relationship between the allosteric effect as measured in McCormick et al. 2024 and the growth rate difference reported in McCormick et al. 2021.

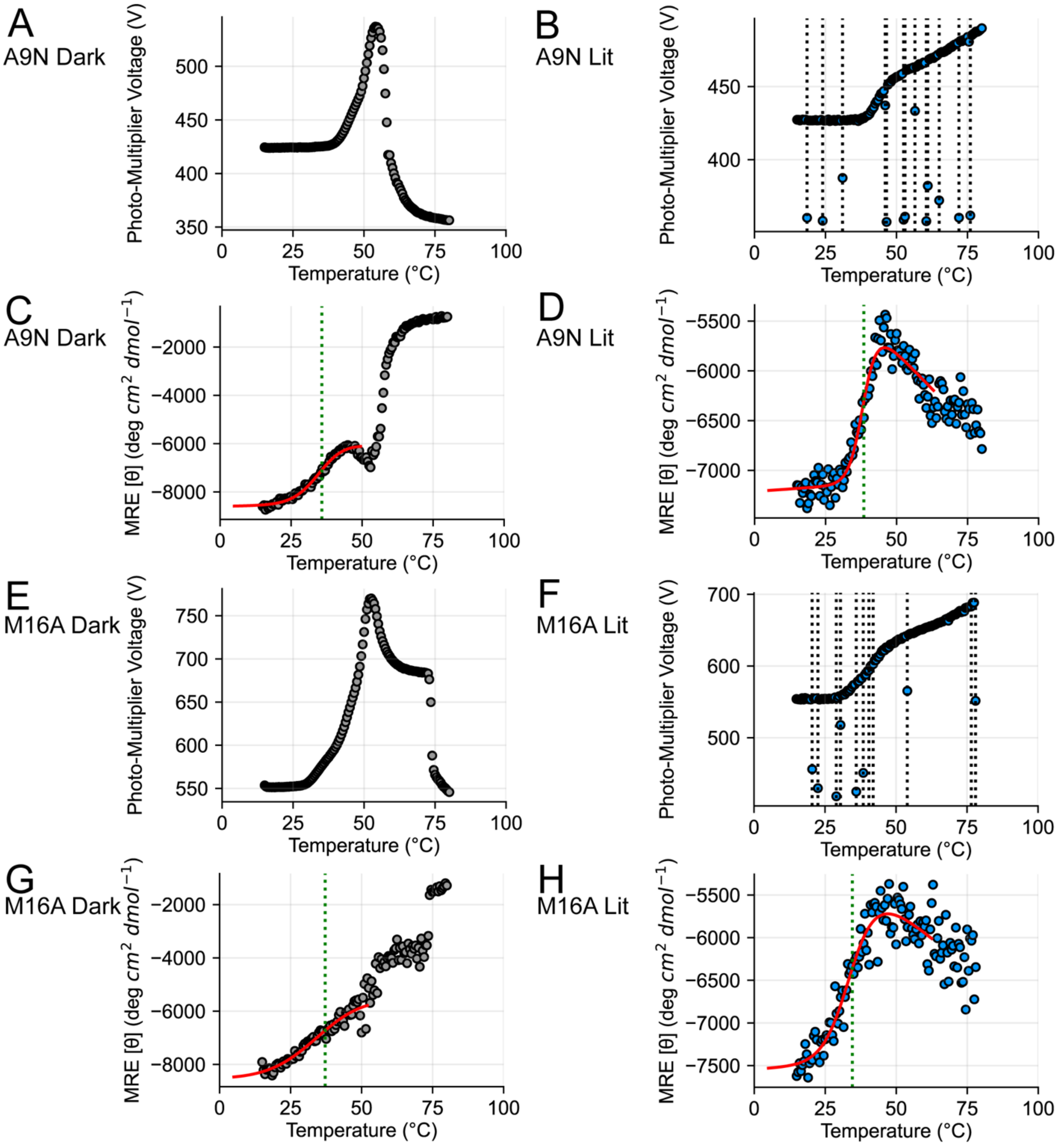


**Figure S7: Thermal melts and associated HT voltage signal for DL121 A9N and DL121 M16A.** The panel layouts, color coding, and markings follow from figure S5.

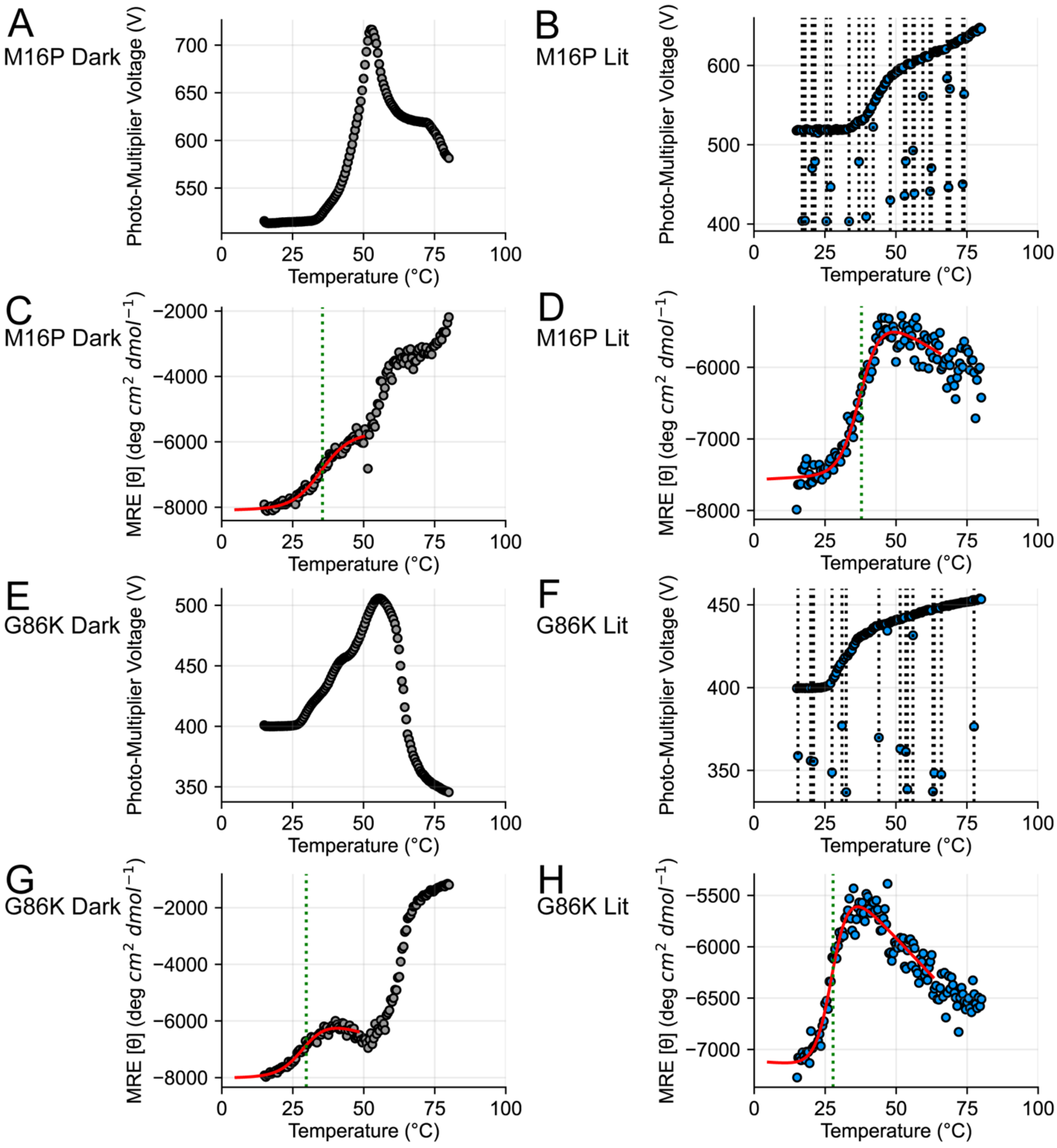


**Figure S8: Thermal melts and associated HT voltage signal for DL121 M16P and DL121 G86K.** The panel layouts, color coding, and markings follow from figure S5.

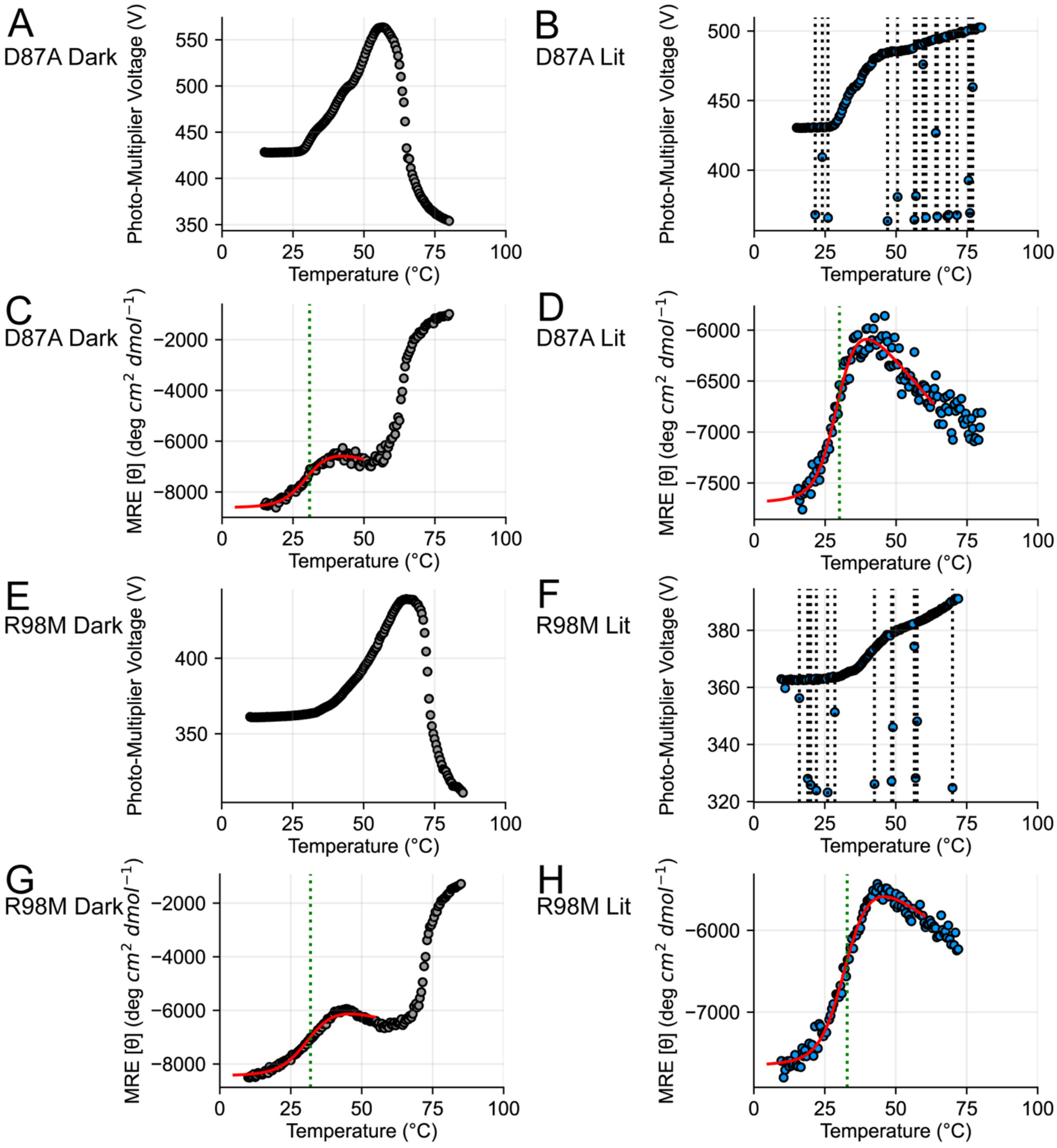


**Figure S9: Thermal melts and associated HT voltage signal for DL121 D87A and DL121 R98M.** The panel layouts, color coding, and markings follow from figure S5.

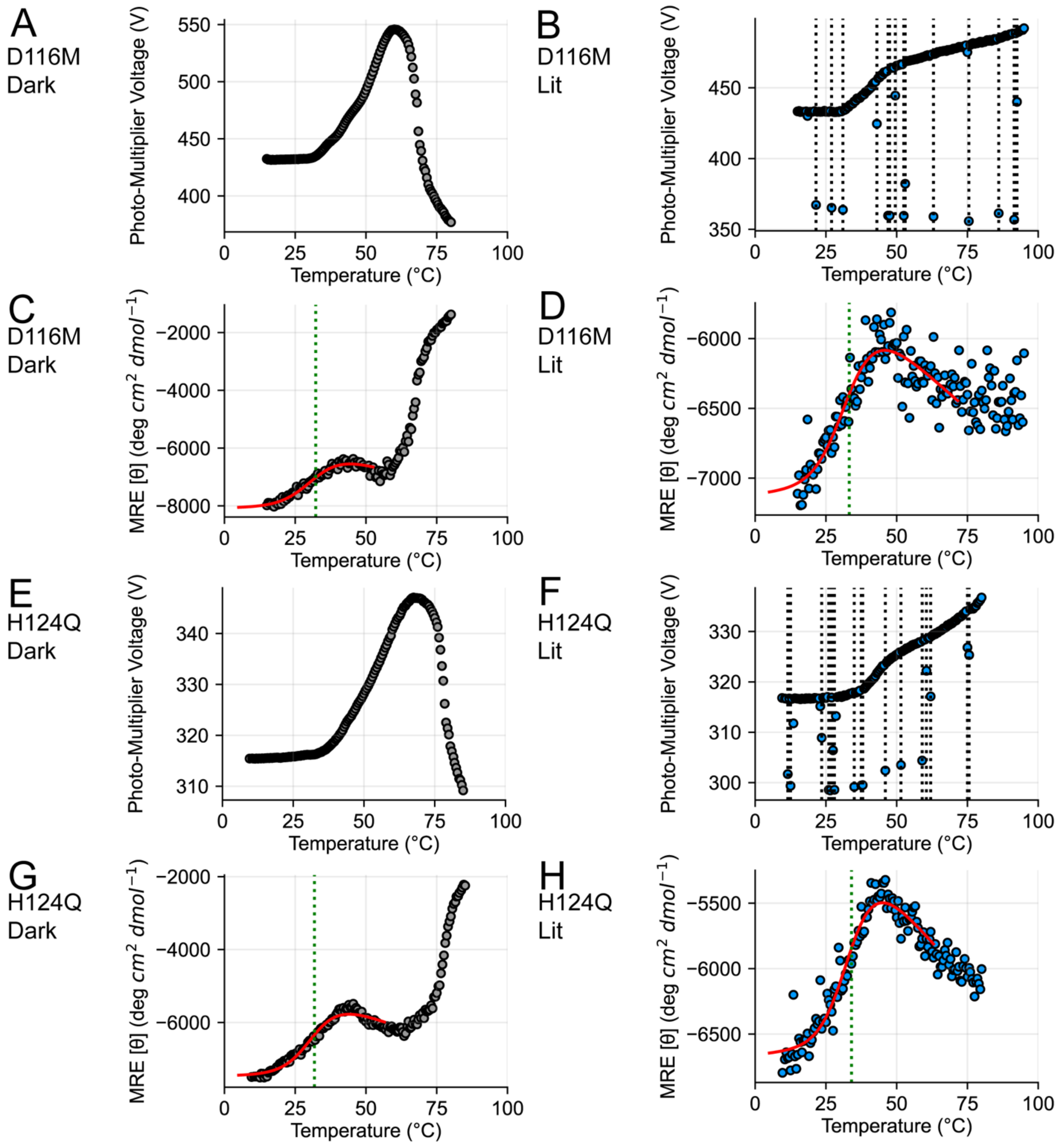


**Figure S10: Thermal melts and associated HT voltage signal for DL121 D116M and DL121 H124Q.** The panel layouts, color coding, and markings follow from figure S5.

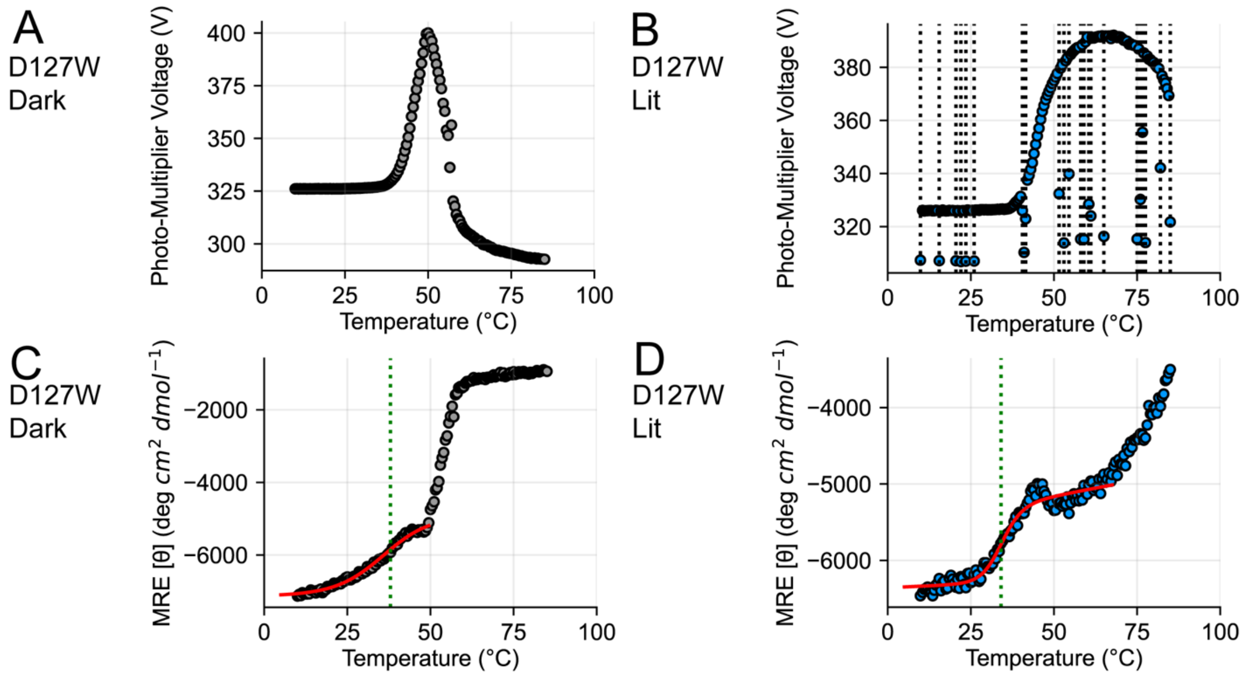


**Figure S11: Thermal melts and associated HT voltage signal for DL121 D127W.** The panel layouts, color coding, and markings follow from figure S5.

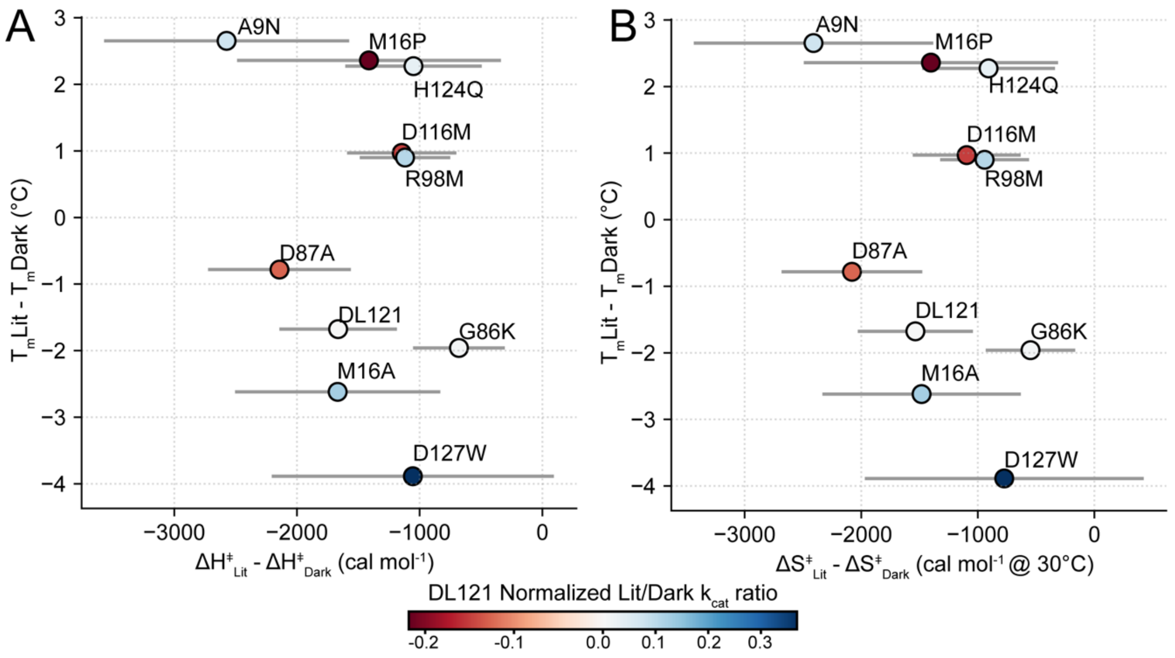


**Figure S12**: **Relationship between thermal stability and transition state enthalpy and entropy.**

A) Relationship between the light-dependent change in the melting point temperature ($T_m$), and the light-dependent change in transition state enthalpy. Each point is color coded by the magnitude of allostery. $T_m$ measurements were made in singlicate. Error bars along the horizontal axis were estimated by bootstrap resampling and refitting the catalytic turnover data shown in Fig. S3 (see methods for details).

B) Relationship between the light-dependent change in the melting point temperature ($T_m$), and the light-dependent change in transition state entropy. Each point is color coded by the magnitude of allostery. $T_m$ measurements were made in singlicate. Error bars along the horizontal axis were estimated by bootstrap resampling and refitting the catalytic turnover data shown in Fig. S3 (see methods for details).

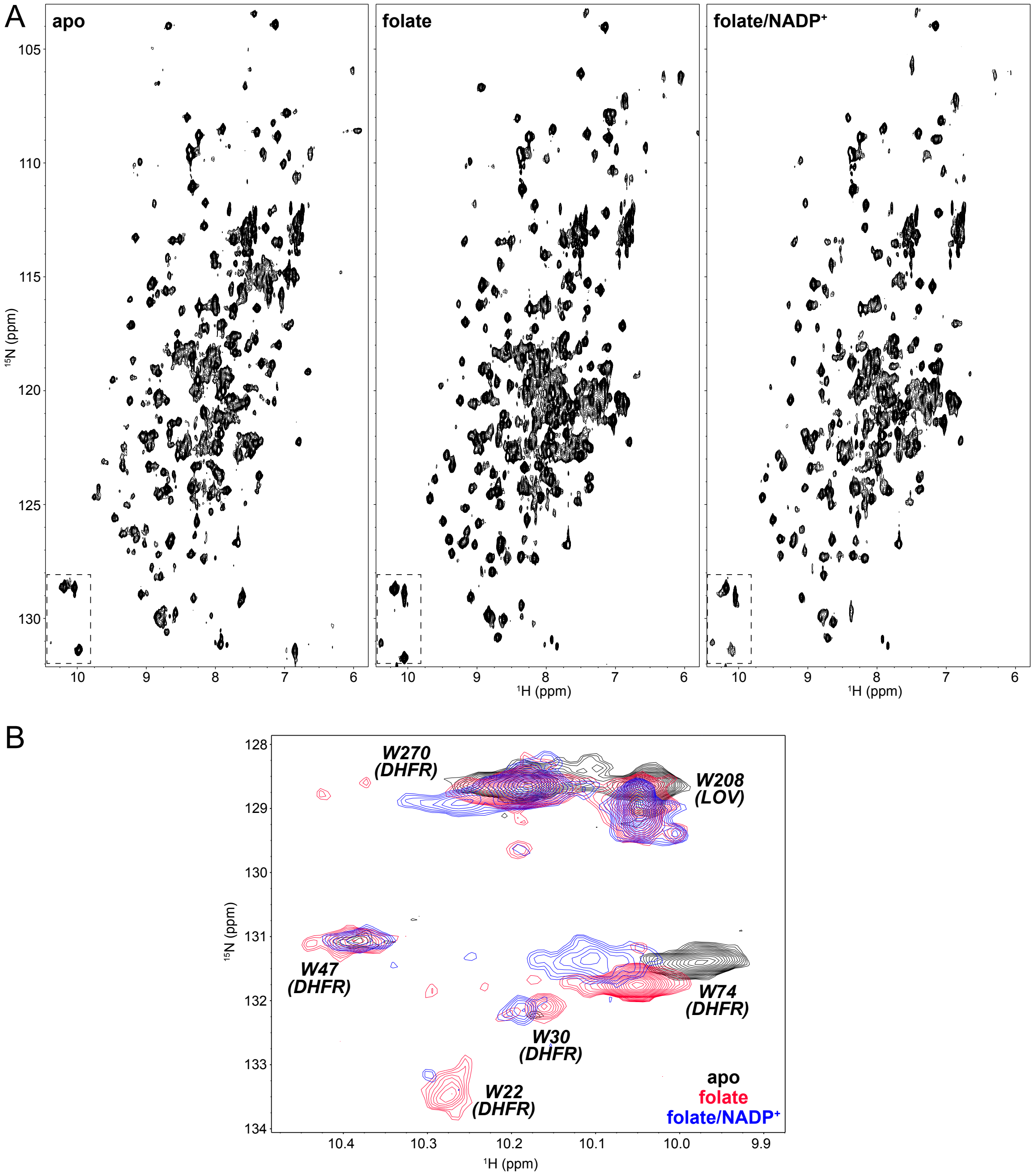


**Figure S13. DL121 NMR chemical shift changes associated with folate binding.**

A) $^{15}N/^{1}H$ TROSY spectra of DL121 either in an apo form or in the presence of 1 mM folate or 1 mM folate and NADP+, as indicated. Dashed lines at bottom left indicate regions used for inset in panel B).

B) Tryptophan indole Hε1/Nε1 region of $^{15}N/^{1}H$ TROSY spectra of DL121 either in an apo form or in the presence of 1 mM folate or 1 mM folate and NADP+, as indicated. Assignments are derived by analogy to previously-reported assignments of LOV2 or DHFR, as indicated in the text.

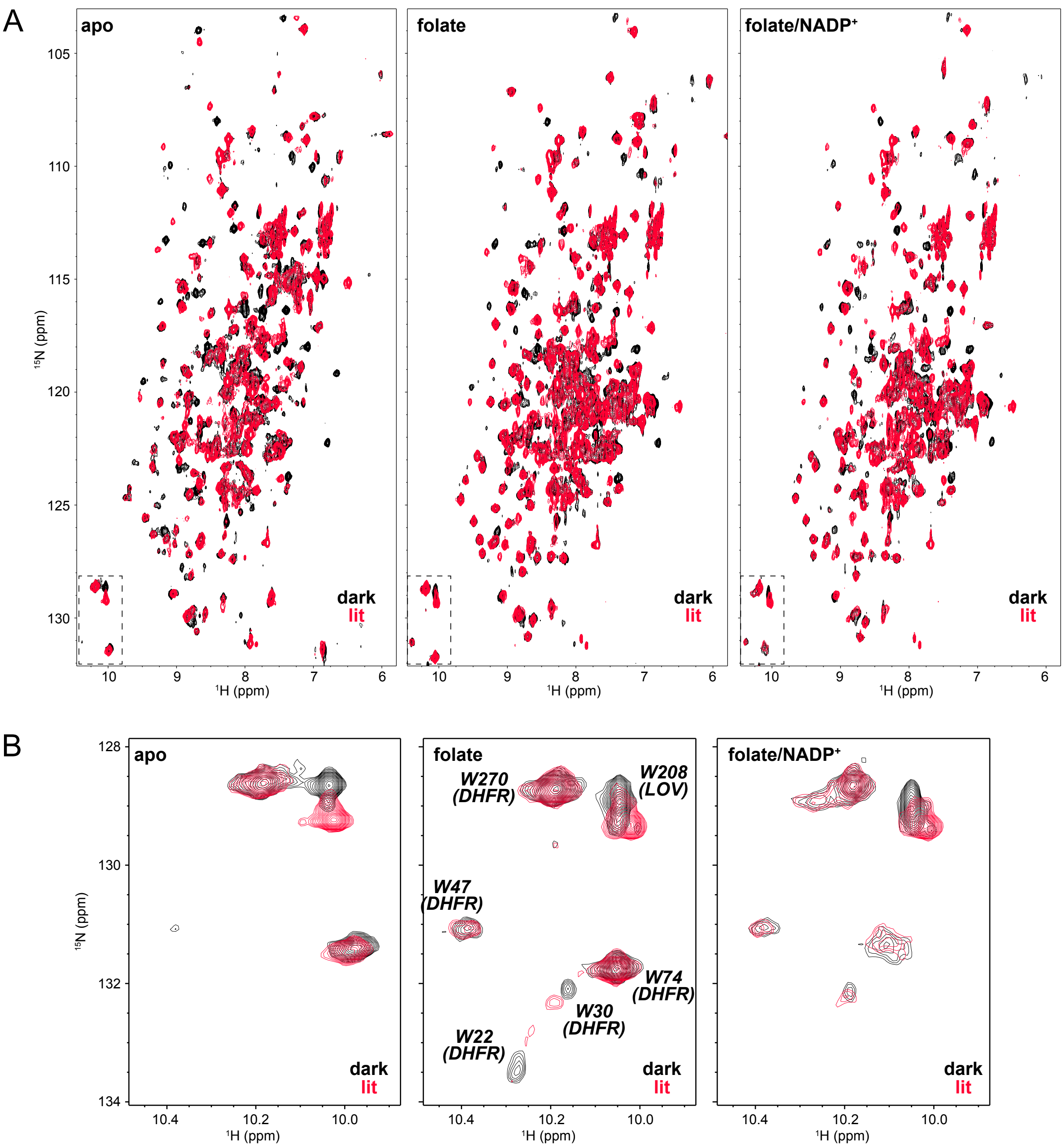


**Figure S14. Light-induced NMR chemical shift changes in DL121.**

A) $^{15}N/^{1}H$ TROSY spectra of DL121 either in an apo form or in the presence of 1 mM folate or 1 mM folate and NADP+, as indicated. Spectra were recorded in the dark (black) or light (red; each NMR scan preceded by 100 ms, 50 mW pulse of 488 nm laser light). Dashed lines at bottom left of folate spectra indicate region the expanded in panel B.

B) Tryptophan indole Hε1/Nε1 region of $^{15}N/^{1}H$ TROSY spectra of DL121 either in an apo form or in the presence of 1 mM folate or 1 mM folate and NADP+, as indicated.

| | DL121 | A9N | M16A | M16P | G86K | D87A | R98M | D116M | H124Q | D127W |
|---|---|---|---|---|---|---|---|---|---|---|
| **ΔH‡ dark (kcal/mol)** | 12.62 | 10.48 | 12.97 | 13.16 | 8.32 | 11.66 | 8.96 | 6.46 | 8.91 | 8.36 |
| **ΔH‡ dark σ** | 0.16 | 0.50 | 0.70 | 0.65 | 0.26 | 0.39 | 0.27 | 0.33 | 0.31 | 0.95 |
| **ΔS‡ dark (cal/mol/K)** | -17.35 | -23.85 | -17.81 | -20.19 | -32.03 | -20.07 | -28.96 | -36.43 | -29.86 | -32.83 |
| **ΔS‡ dark σ** | 0.55 | 1.71 | 2.33 | 2.16 | 0.87 | 1.34 | 0.90 | 1.12 | 1.06 | 3.25 |
| **ΔH‡ lit (kcal/mol)** | 10.95 | 7.91 | 11.30 | 11.74 | 7.64 | 9.52 | 7.84 | 5.31 | 7.86 | 7.30 |
| **ΔH‡ lit σ** | 0.45 | 0.87 | 0.47 | 0.84 | 0.28 | 0.43 | 0.25 | 0.30 | 0.46 | 0.63 |
| **ΔS‡ lit (cal/mol/K)** | -22.42 | -31.80 | -22.70 | -24.81 | -33.84 | -26.93 | -32.07 | -40.05 | -32.86 | -35.38 |
| **ΔS‡ lit σ** | 1.55 | 2.93 | 1.61 | 2.81 | 0.93 | 1.49 | 0.87 | 1.04 | 1.55 | 2.20 |
| **kcat dark (sec-1 at 30 °C)** | 0.81 | 1.06 | 0.36 | 0.08 | 0.63 | 1.01 | 1.02 | 1.51 | 0.70 | 0.40 |
| **kcat dark σ** | 0.010 | 0.049 | 0.010 | 0.003 | 0.013 | 0.028 | 0.017 | 0.036 | 0.016 | 0.029 |
| **kcat lit (sec-1 at 30 °C)** | 1.00 | 1.40 | 0.49 | 0.08 | 0.78 | 1.12 | 1.37 | 1.65 | 0.89 | 0.63 |
| **kcat lit σ** | 0.031 | 0.081 | 0.015 | 0.003 | 0.016 | 0.043 | 0.028 | 0.047 | 0.020 | 0.041 |
| **ΔG‡ dark (cal/mol at 30 °C)** | 17874.20 | 17711.12 | 18368.10 | 19272.36 | 18028.23 | 17742.51 | 17736.47 | 17497.69 | 17957.52 | 18307.09 |
| **ΔG‡ dark σ** | 7.43 | 27.76 | 17.16 | 19.10 | 12.87 | 16.50 | 10.12 | 14.45 | 13.42 | 43.44 |
| **ΔG‡ lit (cal/mol at 30 °C)** | 17745.11 | 17546.98 | 18182.09 | 19261.13 | 17895.83 | 17679.25 | 17558.70 | 17447.13 | 17814.58 | 18024.60 |
| **ΔG‡ lit σ** | 18.31 | 34.49 | 18.57 | 20.59 | 12.31 | 22.73 | 12.22 | 16.94 | 13.45 | 38.72 |
| **ΔΔG‡ (cal/mol at 30 °C)** | -129.09 | -164.14 | -186.01 | -11.22 | -132.40 | -63.26 | -177.77 | -50.56 | -142.94 | -282.49 |
| **ΔΔG‡ σ** | 19.75 | 44.28 | 25.14 | 28.01 | 17.92 | 28.06 | 15.89 | 22.28 | 18.98 | 58.59 |
| **ΔΔS‡*T(cal/mol at 30 °C)** | -1535.53 | -2409.44 | -1482.26 | -1402.10 | -547.62 | -2079.45 | -941.38 | -1095.40 | -907.53 | -772.89 |
| **ΔΔS‡*T(cal/mol at 30 °C)** | 493.76 | 1026.99 | 850.82 | 1090.42 | 383.56 | 603.16 | 380.75 | 462.85 | 569.28 | 1196.18 |
| **ΔΔH‡ (cal/mol)** | -1664.63 | -2573.58 | -1668.27 | -1413.32 | -680.01 | -2142.71 | -1119.16 | -1145.95 | -1050.47 | -1055.38 |
| **ΔΔH‡ (cal/mol) σ** | 478.91 | 998.95 | 835.52 | 1075.05 | 373.22 | 581.71 | 368.72 | 444.74 | 556.34 | 1149.55 |

**Table S1: Transition state thermodynamic quantities and catalytic turnover measurements for DL121 and select allostery tuning mutations**. These parameters resulted from fitting the Eyring equation to the data in figure S3. Error (**σ**) for all quantities was estimated by bootstrap resampling the experimental data and refitting the model 5000 times (see methods for details).

| Mutant | Lit state | Tm (°C) | Tm σ | coefficient of determination | ΔH | ΔH σ | F (MRE) | F σ | U (MRE) | U σ | Cf (correction folded) | Cu (correction unfolded) |
|---|---|---|---|---|---|---|---|---|---|---|---|---|
| DL121 | Dark | 35.14 | 0.03 | 0.98 | -33487.43 | 145.30 | -4663.73 | 36.75 | -8131.82 | 4.76 | 2.00 | -23.29 |
| DL121 | Lit | 33.46 | 0.01 | 0.95 | -52041.63 | 74.72 | -4111.99 | 3.48 | -7148.53 | 2.29 | 2.00 | -29.16 |
| D87A | Dark | 30.83 | 0.01 | 0.98 | -42280.95 | 104.67 | -5011.92 | 11.23 | -8611.95 | 4.95 | 2.00 | -33.63 |
| D87A | Lit | 30.04 | 0.01 | 0.96 | -48740.59 | 66.57 | -4687.50 | 1.87 | -7688.73 | 3.56 | 2.00 | -32.17 |
| M16P | Dark | 35.50 | 0.03 | 0.98 | -36887.44 | 137.29 | -5573.91 | 32.60 | -8088.65 | 3.75 | 2.00 | -2.33 |
| M16P | Lit | 37.86 | 0.00 | 0.97 | -52991.05 | 50.79 | -4224.62 | 2.75 | -7569.04 | 1.40 | 2.00 | -24.12 |
| M16A | Dark | 37.19 | 0.13 | 0.92 | -20425.89 | 246.41 | -3982.16 | 150.86 | -8553.71 | 6.71 | -2.00 | -22.50 |
| M16A | Lit | 34.57 | 0.01 | 0.93 | -39462.73 | 61.94 | -4295.99 | 4.26 | -7543.03 | 2.76 | 2.00 | -27.32 |
| DL116 | Dark | 39.44 | 0.02 | 0.70 | -88373.62 | 468.84 | -5086.19 | 2.34 | -6835.19 | 3.95 | 2.00 | -31.71 |
| DL116 | Lit | 37.54 | 0.01 | 0.35 | -150000.00 | 1242.86 | -5737.76 | 1.13 | -6444.85 | 3.06 | -1.02 | -13.21 |
| A9N | Dark | 35.84 | 0.02 | 0.99 | -39730.42 | 106.95 | -4846.36 | 30.11 | -8599.11 | 3.33 | 2.00 | -22.53 |
| A9N | Lit | 38.49 | 0.00 | 0.95 | -77708.53 | 87.97 | -4347.79 | 2.26 | -7211.53 | 0.99 | 2.00 | -29.34 |
| G86K | Dark | 29.75 | 0.02 | 0.97 | -42309.97 | 131.89 | -4797.20 | 12.13 | -8012.84 | 5.99 | 2.00 | -32.34 |
| G86K | Lit | 27.79 | 0.01 | 0.95 | -60899.78 | 77.62 | -4454.11 | 1.21 | -7113.70 | 3.88 | -2.00 | -29.24 |
| D116M | Dark | 32.25 | 0.03 | 0.97 | -33838.82 | 139.70 | -5098.80 | 15.28 | -8072.25 | 5.59 | 2.00 | -28.84 |
| D116M | Lit | 33.22 | 0.04 | 0.85 | -33259.09 | 80.21 | -5141.25 | 2.66 | -7116.39 | 3.51 | 2.00 | -18.22 |
| C450S | Dark | 34.40 | 0.01 | 0.98 | -57428.60 | 90.57 | -3924.46 | 3.66 | -6938.11 | 2.44 | 2.00 | -23.73 |
| C450S | Lit | 34.00 | 0.01 | 0.98 | -48988.72 | 87.15 | -3681.61 | 4.17 | -6665.19 | 3.02 | 2.00 | -22.34 |
| R98M | Dark | 31.96 | 0.01 | 0.99 | -37985.06 | 57.30 | -4757.24 | 7.14 | -8430.92 | 1.64 | 2.00 | -26.84 |
| R98M | Lit | 32.86 | 0.01 | 0.99 | -41982.39 | 52.30 | -4345.03 | 3.94 | -7646.50 | 1.32 | 2.00 | -24.51 |
| D127W | Dark | 37.96 | 0.15 | 0.99 | -25846.43 | 165.94 | -4193.90 | 92.87 | -7126.98 | 2.06 | 2.00 | -11.88 |
| D127W | Lit | 34.08 | 0.01 | 0.97 | -61182.64 | 108.46 | -5572.46 | 1.48 | -6356.26 | 0.79 | 2.00 | 8.31 |
| H124Q | Dark | 31.77 | 0.01 | 0.97 | -37034.57 | 67.22 | -4390.80 | 5.68 | -7463.68 | 1.65 | 2.00 | -27.99 |
| H124Q | Lit | 34.04 | 0.01 | 0.94 | -37128.64 | 73.21 | -4257.28 | 3.68 | -6658.41 | 1.32 | 2.00 | -24.60 |

**Table S2: CD-derived thermal stability parameters for DL121 and select allostery tuning mutations**. These parameters resulted from fitting of a two state unfolding model to the data in figures S5, S7-S11.
